# *S. pombe wtf* genes use dual transcriptional regulation and selective protein exclusion from spores to cause meiotic drive

**DOI:** 10.1101/2021.09.30.462505

**Authors:** Nicole L. Nuckolls, Ananya Nidamangala Srinivasa, Anthony C. Mok, María Angélica Bravo Núñez, Jeffrey J. Lange, Todd J. Gallagher, Chris W. Seidel, Sarah E. Zanders

**Affiliations:** Stowers Institute for Medical Research, Kansas City, United States; Department of Molecular and Integrative Physiology, University of Kansas Medical Center, Kansas City, United States; University of Missouri—Kansas City, Kansas City, United States

## Abstract

Meiotic drivers bias gametogenesis to ensure their transmission into more than half the offspring of a heterozygote. In *Schizosaccharomyces pombe*, *wtf* meiotic drivers destroy the meiotic products (spores) that do not inherit the driver from a heterozygote, thereby reducing fertility. *wtf* drivers encode both a Wtf^poison^ protein and a Wtf^antidote^ protein using alternative transcriptional start sites. Here, we analyze how the expression and localization of the Wtf proteins are regulated to achieve drive. We show that transcriptional timing and selective protein exclusion from developing spores ensure that all spores are exposed to Wtf4^poison^, but only the spores that inherit *wtf4* receive a dose of Wtf4^antidote^ sufficient for survival. In addition, we show that the Mei4 transcription factor, a master regulator of meiosis, controls the expression of the *wtf4^poison^* transcript. This dual transcriptional regulation, which includes the use of a critical meiotic transcription factor, likely complicates the universal suppression of *wtf* genes without concomitantly disrupting spore viability. We propose that these features contribute to the evolutionary success of the *wtf* drivers.

**Author Summary:** Killer meiotic drivers are one type of selfish DNA sequence. When only one copy of a killer meiotic driver is found in a genome, the driver is expected to be transmitted to only half of the gametes (e.g. eggs or sperm). Killer meiotic drivers, however, kill developing gametes that do not inherit them, giving the driver a transmission advantage into the next generation. The molecular mechanisms used by these killers are not well understood. In this work, we analyzed how one killer meiotic driver, *wtf4* from fission yeast, ensures targeted gamete (spore) killing. Previous work showed that *wtf* meiotic drivers encode a poison protein that is transmitted to all spores and an antidote protein that rescues only spores that inherit the locus. Here, we show that different timing of the expression of the two proteins, combined with differential inclusion of the proteins in developing spores, both contribute to targeted spore killing. We also demonstrate that *wtf4* exploits an essential gene expression pathway, making it difficult for the genome to prevent this locus from being expressed and killing. This extends our knowledge both of how these genetic parasites act and how they are equipped to evade host suppression mechanisms.

## Introduction

The transmission of most eukaryotic genes follows Mendel’s first law of segregation. This law stipulates that the two alleles of a heterozygous organism (e.g. *A/a*) segregate randomly into gametes such that each allele is transmitted to 50% of the progeny (*Fairbanks and Abbot, 2016*). There are, however, alleles that can break Mendel’s law to force their own transmission into more than half of the offspring. These lawbreaking genes are called meiotic drivers (*Burt and Trivers, 2006*; *Lindholm et al., 2016*). There is a tremendous diversity of meiotic drive genes with distinct evolutionary origins and mechanisms found throughout eukaryotes (*Akera et al., 2017; Bauer et al., 2012; Bauer et al., 2007; Chen et al., 2008; Dalstra et al., 2005; Dawe et al., 2018; Didion et al., 2015; Grognet et al., 2014; Hammond et al., 2012; Helleu et al., 2014; Herrmann et al., 1999; Kruger et al., 2019; Larracuente and Presgraves, 2012; Lin et al., 2018; Long et al., 2008; Phadnis and Orr, 2009; Pieper et al., 2018; Rathje et al., 2019*; *Rhoades et al., 2019; Shen et al.,2017; Svedberg et al., 2021; Vogan et al., 2019; Wu et al., 1988; Xie, et al., 2019; Yu, et al., 2018).* However, the molecular details underlying how these systems are expressed and function are limited. Uncovering these details is important for understanding meiotic drive and, more broadly, has the potential to reveal novel insights about gametogenesis. For example, the existence of a sperm-autonomous phenotype, despite cytoplasmic connections between sister sperm, was discovered through study of the *t*-haplotype driver in mouse (*Bauer et al., 2012; Bauer et al., 2007*).

Meiotic drivers can generally be considered selfish or parasitic genes (*Sandler and Novitski, 1957*). This is because drivers can persist in genomes due to the transmission advantages of drive, rather than due to fitness benefits they provide to the organisms that carry them. In fact, meiotic drivers often cause decreased fitness through a variety of direct and indirect mechanisms (*Johnson, 2010; Presgraves, 2010; Ségurel et al., 2011; Zanders and Unckless, 2019*). The fitness costs are especially deleterious amongst the class of drivers known as the killer meiotic drivers (reviewed in *Bravo Núñez et al., 2018b*). These drivers can achieve up to 100% transmission to viable gametes by destroying the products of meiosis that do not inherit the driver from a heterozygote.

Despite the fitness costs of killer meiotic drivers, many populations harbor these genes (reviewed in *Bravo Núñez et al., 2018b*). However, the fission yeast *S. pombe* may represent an extreme case. Different natural isolates of *S. pombe* contain between 4-14 predicted killer meiotic drivers from the *wtf* (<u>w</u>ith transposon fission yeast) gene family (*Bravo Núñez et al., 2020a; Eickbush et al., 2019; Hu et al., 2017; Nuckolls et al., 2017*). These genes kill the meiotic products (spores) that do not inherit them from a heterozygote using two proteins produced from largely overlapping coding sequences: a Wtf^poison^ protein that kills spores and a Wtf^antidote^ protein that rescues spores that inherit the driver (Figure 1B; *Nuckolls et al., 2017*; *Hu et al., 2017).* Largely because of these drivers, heterozygous *S. pombe* diploids generated by crossing distinct isolates generally produce very few viable spores (*Avelar, Perfeito, Gordo, & Ferreira, 2013; Bravo Núñez et al., 2020b; Hu et al., 2015; Jeffares et al., 2017; Singh and Klar, 2002; Zanders et al., 2014*). Moreover, the *wtf* genes confer no known fitness benefits, yet are present in multiple copies in all sequenced isolates of *S. pombe* (*Eickbush et al., 2019*; *Hu et al., 2017*).

**Figure 1.**
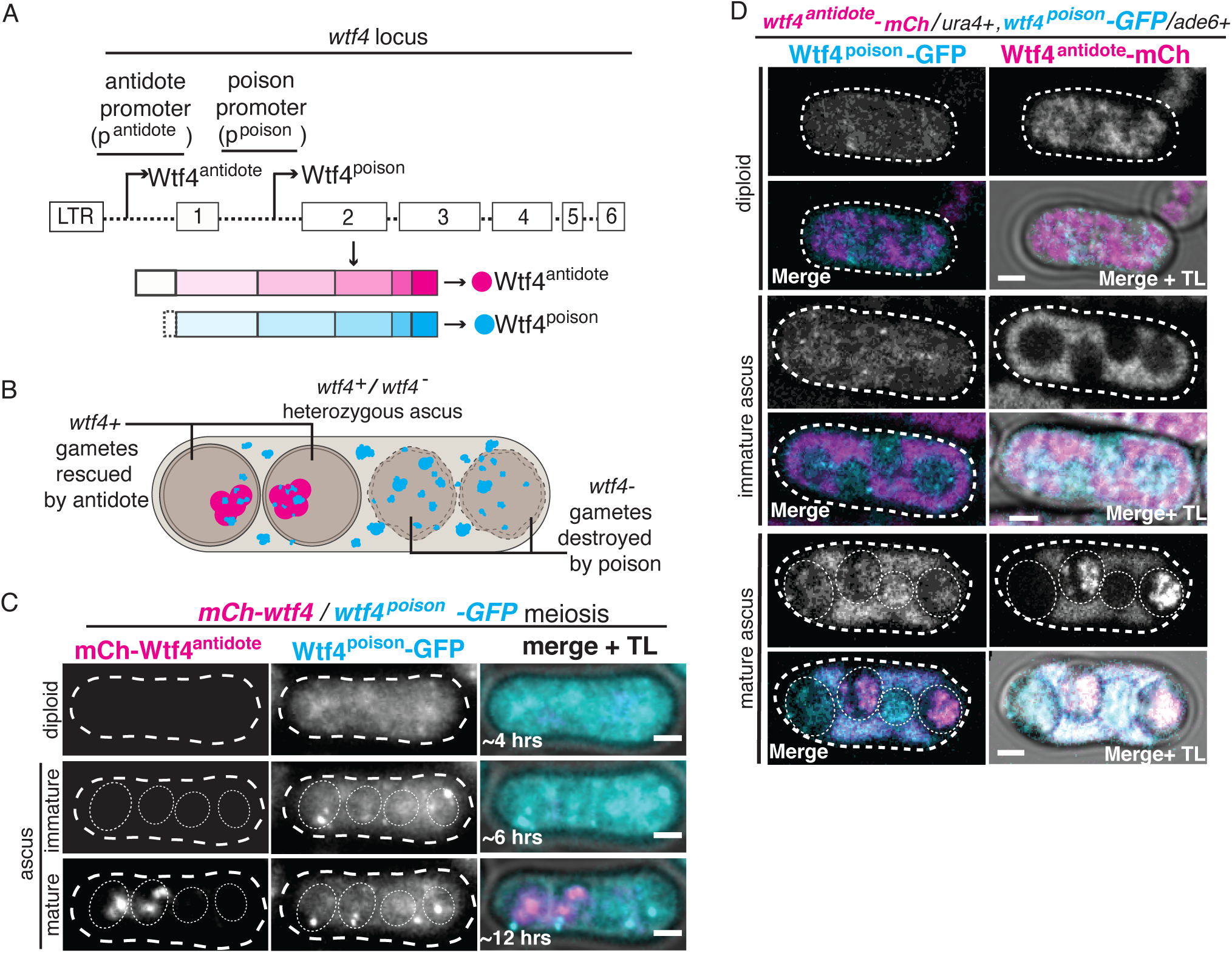
C-terminal tag reveals Wtf4^antidote^ expression prior to spore formation. (**A**) A depiction of the *wtf4* gene is shown. There is a Tf retrotransposon LTR located 284 base pairs upstream of exon 1. The sequence upstream of exon 1 contains the antidote promoter (*p^antidote^*) and intron 1 contains the poison promoter (*p^poison^*). The arrows represent the predicated transcriptional start sites where the majority of the *wtf* transcripts begin (*Kuang et al., 2017*). Wtf4^antidote^ (magenta circle) is encoded by exons 1-6. Wtf4^poison^ (cyan circle) is encoded by exons 2-6. (**B**) Model of Wtf4 poison-antidote meiotic drive from *wtf4* heterozygotes. Wtf4^poison^ is present in all four spores, while Wtf4^antidote^ expression is enriched in only two of the four spores. (**C**) Time-lapse microscopy of a *mCherry-wtf4/wtf4^poison^-GFP* diploid undergoing meiosis and sporulation. mCherry-Wtf4 is shown in magenta and Wtf4^poison^-GFP is shown in cyan in merged images. Images shown at ∼ 4 hours (diploid), ∼8 hours (immature ascus), and ∼12 hours (mature ascus) post-induction in sporulation media. Distinct images for these strains were first presented in (*Nuckolls et al., 2017*) and a distinct meiotic timelapse of these strains was also presented in (*Nuckolls et al., 2020*). (**D**) Images of a heterozygous *wtf4^antidote^-mCherry/ura+, wtf4^poison^-GFP/ade6+* diploid cell, an immature ascus, and a mature ascus. Wtf4^antidote^-mCherry is shown in magenta and Wtf4^poison^-GFP is shown in cyan in merged images. Images of this cross are also depicted in Figure 4D. Images were taken after 3 days on sporulation media. TL= transmitted light. All scale bars represent 2 μm. Not all images are shown at the same brightness and contrast to avoid over saturation of pixels in the brighter images.

Due to the fitness costs that *wtf* drivers cause during meiosis, *S. pombe* has likely evolved ways to tolerate or suppress these drivers. There are currently two described mechanisms that can suppress *wtf* meiotic drivers: suppression by other *wtf* genes and gamete disomy (*Bravo Núñez et al., 2018a; Bravo Núñez et al., 2020b*). Transcriptional silencing acting through trans-acting regulators or cis-acting chromatin packaging is an additional candidate suppression mechanism. Indeed, transcriptional silencing is critical for suppressing transposons, the most widespread and thoroughly studied selfish genetic elements (*McClintock, 1951; Sienski et al., 2012; Slotkin and Martienssen, 2007*). On the chromatin side, histone deacetylation and TOR-mediated RNA processing have been shown to contribute to the mitotic silencing of both Tf2 retrotransposons and the *wtf* genes in *S. pombe* (*Hansen et al., 2005; Nicolas et al., 2007; Watts et al., 2018; Wei et al., 2021*).

To find possible routes to *wtf* driver suppression through transcriptional regulation, we sought to further characterize the transcriptional regulation of the *wtf* drivers using the *wtf4* gene as a model. We define the promoter sequences that control the *wtf4^poison^* and *wtf4^antidote^* transcripts. We demonstrate that staggered expression due to the distinct promoters and differential inclusion of the two Wtf4 proteins in developing spores both contribute to Wtf4 protein localization and thus efficient drive. We also found that the expression of the *wtf4^poison^* transcript is controlled by the Mei4 transcription factor, a master regulator of gene expression in meiosis (*Alves-Rodrigues et al., 2016; Ioannoni, et al., 2016;*). In the context of our results, we discuss why evolving or maintaining transcriptional silencing of *wtf* genes and drivers in general may be challenging.

## Results

### C-terminal tag reveals an additional population of Wtf4^antidote^ protein

Here, we use the *wtf4* allele from a natural isolate of *S. pombe* known *as S. kambucha,* as a case-study to understand how the expression and localization of Wtf driver proteins is coordinated to ensure meiotic drive. We test the *S. kambucha wtf4* allele within the lab isolate of *S. pombe,* as we previously showed that *wtf4* exhibits strong drive when heterozygous (*Nuckolls et al., 2017*). Previous work demonstrated that *wtf* drivers encode Wtf^poison^ and Wtf^antidote^ proteins using two distinct transcripts. The *wtf^antidote^* transcript is longer and starts upstream of exon 1. The shorter *wtf^poison^* transcript begins upstream of exon 2, within intron 1 of the *wtf^antidote^* transcript (*Eickbush et al., 2019; Hu et al., 2017; Kuang et al., 2017*; *Nuckolls et al., 2017*; Fig 1A).

In previous work, we used fluorescent markers to separately tag and visualize the Wtf4 proteins (*Nuckolls et al., 2017, Nuckolls et al., 2020*). In that work, we found that the Wtf4^poison^-GFP protein is present at low levels in diploid cells induced to undergo meiosis. After meiosis, we observed higher levels of the Wtf4^poison^-GFP protein within both the spores and the ascus (sac) that holds the spores (*Nuckolls et al., 2017*). In contrast, we detected mCherry-Wtf4^antidote^ protein strongly enriched within spores that inherited the tagged *wtf4* allele (*Nuckolls et al., 2017).* These distribution patterns explain how the Wtf4^poison^ protein has the potential to kill all spores and shows how the Wtf4^antidote^ specifically rescues the spores that inherit the *wtf4* allele (Fig 1B). This expression pattern did, however, raise the question as to how meiotic cells survived expression of the Wtf4^poison^ prior to strong expression of the Wtf4^antidote^ in spores. This led us to question if the mCherry-Wtf4^antidote^ tag revealed the full spectrum of Wtf4^antidote^ expression.

In this work, we explored the expression of the two Wtf4 proteins more thoroughly to uncover how the two proteins achieve targeted destruction of spores that do not inherit *wtf4* from a heterozygote (Fig 1B). We again used *wtf4^poison^-GFP*, a separation-of-function allele that does not express Wtf4^antidote^ and adds a GFP tag to the C-terminus of the Wtf4^poison^. We have previously shown that this allele acts as a functional poison but is slightly less toxic than the corresponding untagged *wtf4^poison^* allele (*Nuckolls et al., 2017,* S1A Fig, diploid 4). We also used the *mCherry-wtf4* allele, which tags Wtf4^antidote^, but still generates an untagged Wtf4^poison^. This allele functions similarly to the wild-type *wtf4* in allele transmission and viable spore yield (VSY) assays (*Nuckolls et al., 2017,* S1A Fig, diploid 3). VSY is a measure of fertility that assays the number of viable spores generated per cell induced to form spores (*Smith, 2009*). In this work, we normalized viable spore yields to those measured in wild-type control cells with an empty vector integrated in the genome instead of a *wtf4* allele. Those wild-type cells are therefore considered to have 100% fertility. Complete drive in a heterozygote is expected to reduce fertility by roughly 50% and expression of a Wtf4^poison^ allele in the absence of Wtf4^antidote^ is expected to reduce fertility more than 50%.

We integrated the *mCherry-wtf4* and *wtf4^poison^-GFP* alleles mentioned above into the *ade6* locus of different haploid strains. We crossed these two strains together to generate heterozygous *mCherry-wtf4 / wtf4^poison^-GFP* diploids. We then completed time-lapse microscopy during fission yeast meiosis and spore formation (Fig 1C). We depict Wtf4^antidote^ in magenta and Wtf4^poison^ in cyan and this will be true for all images in this text regardless of fluorescent protein tag. This analysis confirmed our previous gross observations of early and late meiotic time points (*Nuckolls et al., 2017; Nuckolls et al., 2020*), in which we observed Wtf4^poison^-GFP prior to the bulk of the mCherry-Wtf4^antidote^ signal. Indeed, we saw Wtf4^poison^-GFP hours before mCherry-Wtf4^antidote^ and given that GFP maturation is only minutes faster than mCherry (*Badrinarayanan et al., 2012; Khmelinskii et al., 2012; Shashkova et al., 2018*), the earlier expression of Wtf4^poison^-GFP cannot be explained by maturation times of the fluorescent proteins alone. We saw similar expression dynamics of the mCherry-Wtf4^antidote^ protein expressed from a separation-of-function allele, *mCherry-wtf4^antidote^*, that does not express the Wtf4^poison^ (S1B Fig and S1A Fig, diploids 8 and 9).

To analyze the expression of the Wtf4^antidote^ more rigorously, we also assayed a C-terminally tagged Wtf4^antidote^ separation-of-function allele, *wtf4^antidote^-mCherry*. This allele does not encode Wtf4^poison^ because it only contains the *wtf4^antidote^-mCherry* coding sequence and not intron 1, which contains the transcriptional and translational start site of the Wtf4^poison^ (Figure 1A). We integrated this allele into the *ura4* locus and found that it indeed acted as an antidote-only allele in that it could suppress drive of *wtf4*, but not cause drive on its own (S1A Fig, diploids 6 and 7). We then crossed a strain carrying the *wtf4^antidote^-mCherry* allele to haploid cells carrying *wtf4^poison^-GFP* to generate *wtf4^antidote^-mCherry/ura4+, wtf4^poison^-GFP/ade6+* diploids. We imaged these diploids and observed that the Wtf4^antidote^ localization was slightly different than the pattern we observed with the *mCherry-wtf4* allele in that the cells had mCherry signal prior to spore formation. Specifically, we saw Wtf4^antidote^-mCherry dispersed throughout diploid cells induced to undergo meiosis and in the ascal cytoplasm of immature asci (Fig 1D, S1C Fig). In mature asci, Wtf4^antidote^-mCherry was mostly found enriched in two of the four spores, similar to the N-terminally tagged protein (Fig 1D). We saw that localization of another C-terminally tagged Wtf4^antidote^ separation-of-function allele, *wtf4^antidote^-GFP*, localized similarly to *wtf4^antidote^-mCherry* (S1D Fig) and behaved as a functional antidote allele (S1A Fig, diploid 11,13).

We conclude that the C-terminal tag reveals the production of a Wtf4 protein population that is not apparent with the N-terminally tagged allele. Although the nature of the additional protein is not clear, it could be produced using the second translational start site found in exon 1 (codon 12). We have previously shown that this start site can be used to encode a fully functional antidote protein (*Nuckolls et al., 2017*). Importantly, our results suggest that at least some Wtf4^antidote^ is present to potentially neutralize the Wtf4^poison^ prior to spore formation. This interpretation is also consistent with previous long-read RNA sequencing data showing at least some transcription of *wtf* antidotes prior to spore formation (e.g. 0-6 hours after meiotic induction (*Kuang et al., 2017*; *Nuckolls et al., 2017; Hu et al., 2017*).

### Distinct promoters are sufficient to explain the distinct localization patterns of the Wtf4 proteins

We next wanted to explore if the localization of the Wtf proteins could be attributed to the transcriptional timing of their respective promoters. To assay this, we generated constructs with fluorescent proteins, mCherry and GFP, under the control of the *wtf4^antidote^* and *wtf4^poison^* promoters, respectively. For the *wtf4^antidote^* promoter (*p^antidote^*), we used the 285 base pairs found upstream of exon 1. This is just downstream of a nearby Tf transposon long terminal repeat (LTR) that is not necessary for Wtf^antidote^ production (Fig 1A; *Hu et al., 2017*; *Nuckolls et al., 2020*). Previous work assaying *cw9* and *cw27* (two *wtf* drivers in the CBS5557 isolate) found that the 288 bp upstream of exon 1 was insufficient to generate a fully functional *p^antidote^* promoter (*Hu et al., 2017*). Although the promoter regions of the genes are highly similar, our previous results showed that for *wtf4,* 285 bp upstream sequence was sufficient to encode a functional Wtf4^antidote^ (*Nuckolls et al., 2020*). Our results suggest that this region includes the promoter of *wtf4^antidote^* and consistent with this idea, this sequence is well conserved amongst *wtf* genes that encode for an antidote (*Bowen et al., 2003*; *Hu et al., 2017; Kuang et al., 2017*; *Nuckolls et al., 2017;* Fig 2A-B). To characterize the poison promoter (*p^poison^*), we used the 230 bp sequence that makes up intron 1 of the antidote transcript (Fig 2C). This sequence includes the transcriptional start site of the *wtf4^poison^* transcript and is well conserved amongst *wtf* drivers (Fig 2C-D, *Eickbush et al., 2019; Hu et al., 2017; Kuang et al., 2017; Nuckolls et al., 2017*).

**Figure 2.**
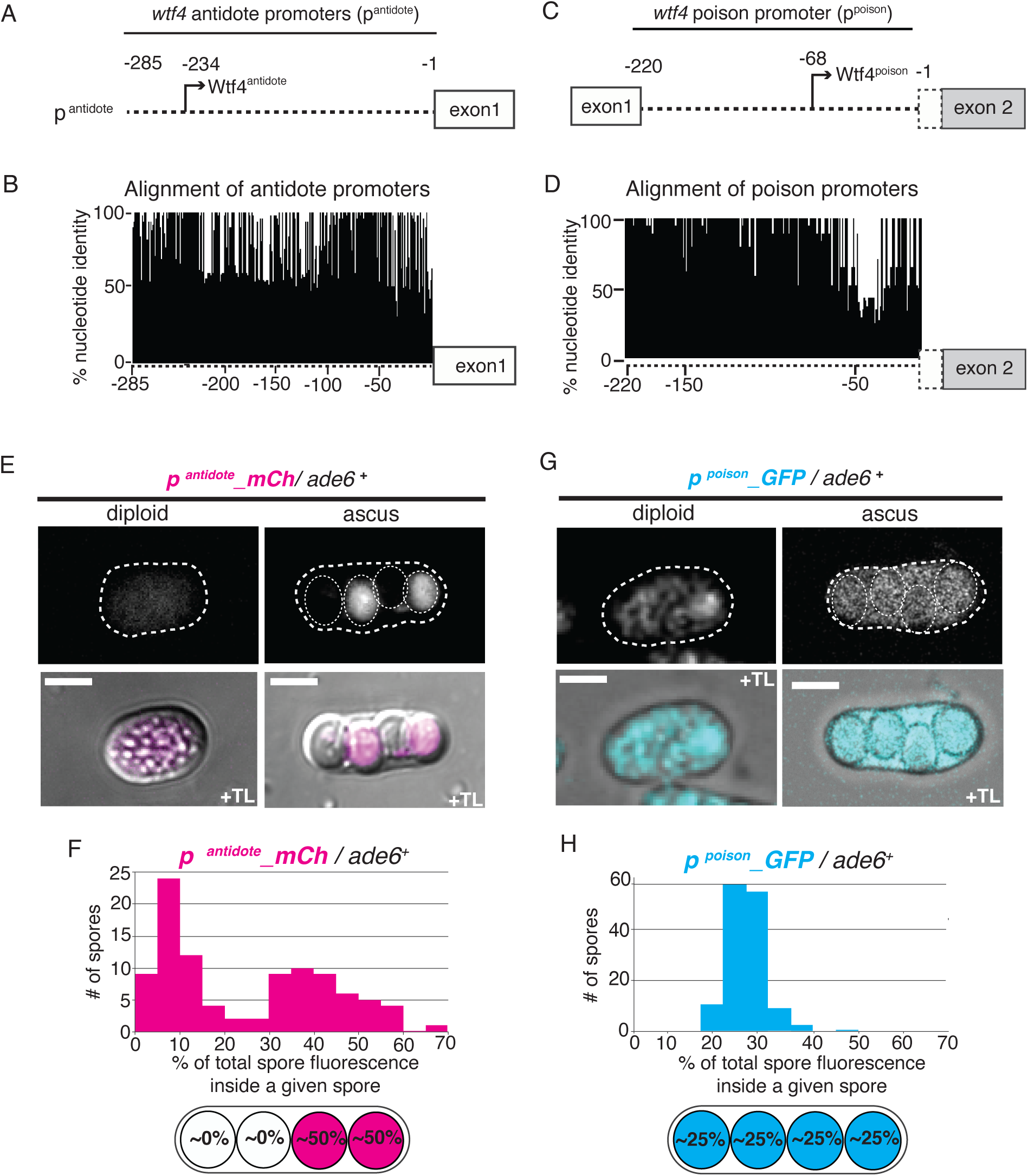
Distinct promoters largely explain differential localization of Wtf4^poison^ and Wtf4^antidote^ proteins. Depictions of (**A**) the *wtf4* antidote promoter, (**B**) the antidote promoter alignments, (**C**) the *wtf4* poison promoter, and (**D**) the poison promoter alignments are shown. For the alignment of antidote promoters, we aligned the 600 base-pairs upstream of 41 predicted antidote-only alleles (*Eickbush et al., 2019*) from three different strains of *S. pombe* (the reference genome*, S. kambucha,* and *FY29033, Lock et al., 2018*). For the alignment of poison promoters, we aligned the intron 1 sequences of 28 predicted poison-antidote *wtf* drivers (*Eickbush et al., 2019*) from three different strains of *S. pombe* (the reference genome*, S. kambucha,* and *FY29033*). The image shows the percent identity at each nucleotide position, excluding gaps. (**E**) Images of *p^antidote^*_*mCherry/ade6+* diploid and ascus. (**F**) Quantification of mCherry fluorescence within *p^antidote^*_*mCherry/ade6+* asci. 24 asci were quantified. (**G**) Images of *p^poison^*_*GFP/ade6+* diploid and ascus. (**H**) Quantification of GFP fluorescence within *p^poison^*_*GFP/ade6+* asci. 34 asci were quantified. All images were acquired after 3 days on sporulation media. For quantification, we assayed the fluorescence intensity within each spore and then divided that number by the total fluorescence intensity within all 4 spores. TL= transmitted light. All scale bars represent 2 μm. Images of diploids are shown at the same brightness and contrast and were imaged at the same settings as the ascus generated from diploids of the same genotype.

We integrated the promoter reporter constructs at the *ade6* locus of different haploid strains. We then generated diploid cells heterozygous for each of the reporters individually (i.e. *reporter*/*ade6*+) and imaged them through meiotic induction and spore formation. With the *p^antidote^_mCherry* reporter, we observed strong mCherry signal in two out of the four spores (Fig 2E-F), and low expression in diploids undergoing meiosis. This suggests the majority of the *p^antidote^_mCherry* reporter is transcribed after the spores individualize. This localization is similar to our observations with tagged Wtf4^antidote^ proteins.

As mentioned above, a fully functional Wtf4^antidote^ protein can be made using a second ATG codon at the 12^th^ codon position (*Nuckolls et al., 2017*). We speculated that additional transcriptional regulatory sequences may be found in those coding sequences upstream of codon 12. To test this, we generated *p^antidote long^_mCherry*, a construct with mCherry under a *wtf4^antidote^* promoter that also contains the first 11 codons of the *wtf4^antidote^* coding sequence (S2A Fig). We integrated this allele at *lys4* and crossed this strain to wild type to generate heterozygous diploids. In these diploids and in the asci generated via these diploids, we saw mCherry expressed from the *p^antidote long^_mCherry* reporter at a similar level to the mCherry expressed from the shorter *p^antidote^* (S2B-C Fig).

For the *p^poison^_GFP* reporter, we observed expression in diploid cells induced to undergo meiosis. We also observed similar GFP reporter signal among the four spores (Fig 2G and 2H). The roughly equal distribution of signal in the four spores, despite the fact that only two inherited the *p^poison^-GFP* reporter, suggests the majority of the reporter transcripts were produced prior to spore individualization. These observations are similar to the localization patterns we observe with Wtf4^poison^-GFP (e.g. Figure 1C).

Together, these experiments demonstrate differential transcription between the *p^antidote^* and *p^poison^* promoters. Our results also show that differential transcription largely explains the localization of Wtf4^poison^ protein within all spores and the enrichment of the Wtf4^antidote^ protein within the spores that inherit the *wtf4* locus.

### Wtf4^antidote^ but not Wtf4^poison^ is excluded from developing spores

The expression of some Wtf4^antidote^ protein prior to spore formation (Fig 1D, S1C Fig, S1D Fig) led us to question why this antidote is insufficient to rescue spores that do not inherit *wtf4+* from *wtf4+*/*wtf-* heterozygotes. A partial answer to this paradox emerged from images of C-terminally tagged *wtf4^antidote^-mCherry* and *wtf4^antidote^-GFP* alleles (Fig 3A, Fig1D, S1C Fig, S1D Fig). These images suggested that the Wtf4^antidote^ made prior to spore formation was excluded from the newly developing spores, as more signal was present in the ascal cytoplasm than in the spores that did not inherit the *wtf4^antidote^-mCherry* allele (Fig 3A, Fig 1D, S1D Fig). We speculated that Wtf4^antidote^ protein made prior to spore formation could protect the cell from the Wtf4^poison^ present during meiosis. But, if the Wtf4^antidote^ protein is excluded from spores and the Wtf4^poison^ is not, the developing spores would need to produce additional Wtf4^antidote^ to protect them from the Wtf4^poison^.

**Figure 3.**
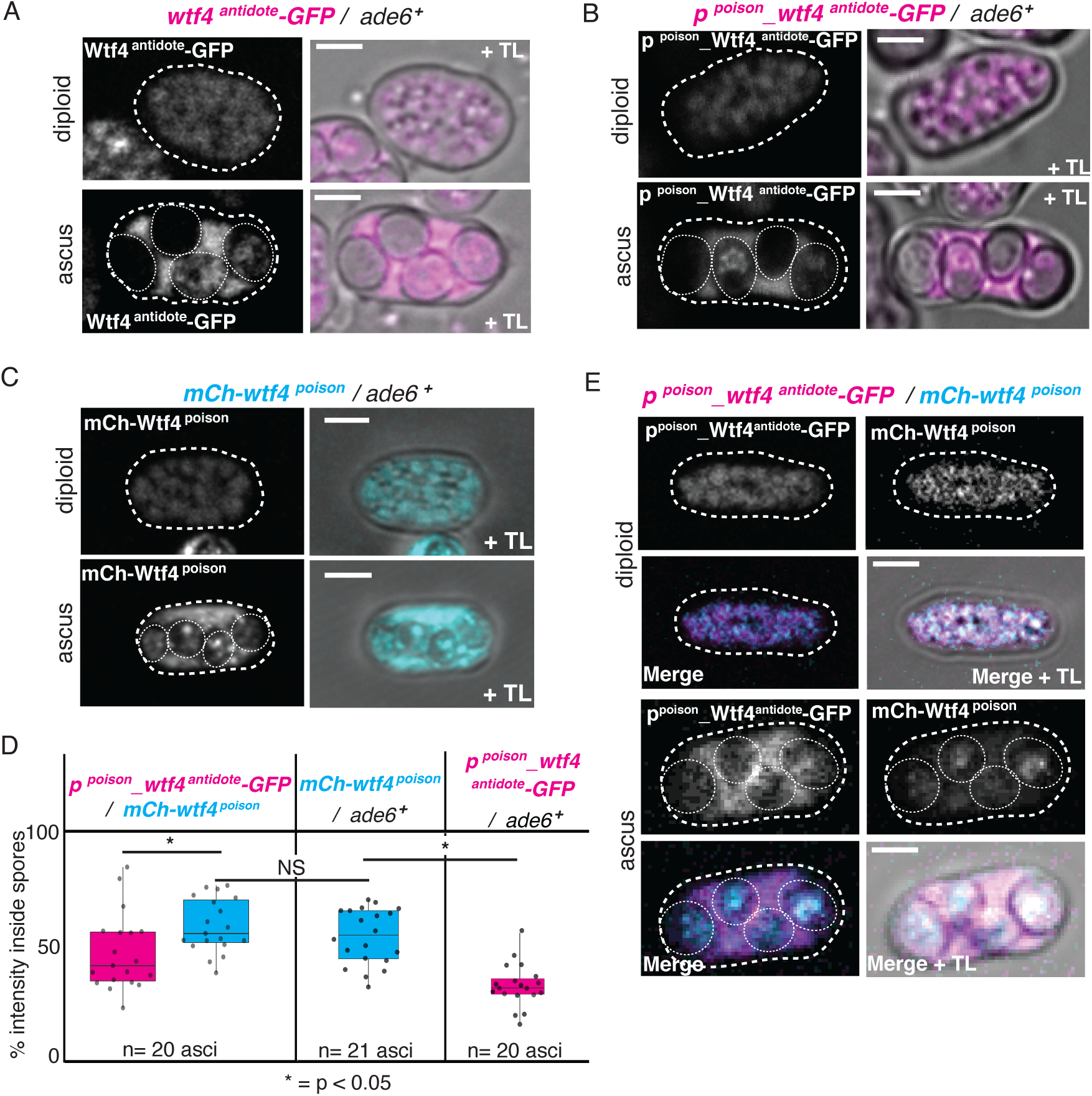
Wtf4^antidote^ is excluded from developing spores more than the Wtf4^poison^. (**A**) Image of a *wtf4^antidote^-GFP/ ade6+* diploid and ascus. Images of this cross are also presented in S1D Fig. (**B**) Image of a *p^poison^*_*wtf4^antidote^-GFP/ ade6+* diploid and ascus. (**C**) Image of a *mCherry-wtf4^poison^/ ade6+* diploid and ascus. (**D**) Quantification of the percentage of the total fluorescence intensity found inside of spores of mature asci produced by *p^poison^_wtf4^antidote^- GFP*/*mCherry-wtf4^poison^* diploids (left, n=20), *mCherry-wtf4^poison^/ade6^+^* diploids (center, n=21) and *p^poison^_wtf4^antidote^-GFP/ade6^+^* diploids (right, n=20). p^poison^_Wtf4^antidote^-GFP fluorescence data is depicted in magenta while mCherry-Wtf4^poison^ is depicted in cyan. Error bars depict standard deviation. NS= not significant. (E) Image of a *p^poison^*_*wtf4^antidote^-GFP/ mCh*-*wtf4^poison^* diploid and ascus. p^poison^_Wtf4^antidote^-GFP is shown in magenta and mCherry-Wtf4^poison^ is shown in cyan in merged images. TL= transmitted light. All scale bars represent 2 μm. All images were acquired after 3 days on sporulation media. Not all images are shown at the same brightness and contrast to avoid over saturation of pixels in the brighter images.

To more thoroughly test the idea that Wtf4^antidote^, but not Wtf4^poison^, is excluded from spores, we wanted to increase the level of Wtf4^antidote^ present at the time of spore formation to a level more comparable to that of the Wtf4^poison^. To do this, we created a construct with Wtf4^antidote^-GFP under the control of the poison promoter (*p^poison^_wtf4^antidote^-GFP*) and generated *p^poison^_wtf4^antidote^-GFP/ade6+* diploids. We then induced the cells to undergo meiosis and imaged them. We observed Wtf4^antidote^-GFP in cells undergoing meiosis (Fig 3B). In mature asci, much of the protein was in the ascal cytoplasm, but there was also Wtf4^antidote^-GFP in two of the four spores (Fig 3B). We speculated that the Wtf4^antidote^-GFP in the two spores is caused by low levels of transcription from the *p^poison^* promoter that occurs after spore individualization. This interpretation is consistent with photobleaching analyses (S3 Fig) and our prior observation that spores that inherit a separation-of-function *wtf4^poison^* allele from a heterozygote are more likely to die than those spores that do not (*Nuckolls et al., 2017*).

We compared the localization of the Wtf4^antidote^-GFP driven by the *p^poison^* promoter to that of Wtf4^poison^ driven by the same promoter to test the hypothesis that the Wtf^antidote^ protein is preferentially excluded from spores. In order to do that, we generated an *mCherry-wtf4^poison^* allele. We integrated the *mCherry-wtf4^poison^* allele at *ade6,* generated *mCherry-wtf4^poison^/ade6+* heterozygous diploids, induced the cells to undergo meiosis, assayed allele transmission, and imaged them. We saw that the allele is indeed functional in that it can cause spore death on its own (S4A Fig, diploid 10). As with our other tagged Wtf4^poison^ constructs, we observed that the mCherry-Wtf4^poison^ protein was more evenly distributed between the spores and the ascal cytoplasm than Wtf4^antidote^-GFP produced from *p^poison^* promoter (Fig 3B and 3C). We quantified how much of each protein made it into spores and found that an average of 34% of the Wtf4^antidote^-GFP fluorescence from the *p^poison^_wtf4^antidote^-GFP* allele was in spores (Fig 3D-right), while the rest remained in the ascal cytoplasm. With mCherry-Wtf4^poison^, however, we observed an average of 55% of the fluorescence inside of spores (Fig 3D-middle). These observations support the idea that Wtf4^antidote^ made prior to spore individualization is excluded from the developing spores at a significantly higher rate than Wtf4^poison^ (p< 0.05, t-test).

We next wanted to test if the presence of the Wtf4^antidote^ in the ascus at the time of spore formation could affect the localization of the Wtf4^poison^ protein. We previously observed in vegetative cells that the Wtf4^antidote^ could promote the localization of the Wtf4^poison^ to the vacuole. We also observed colocalization of Wtf4^antidote^ and Wtf4^poison^ proteins in vacuoles of developing spores (*Nuckolls et al., 2020*). Because vacuoles found in the diploid cell remain in the ascal cytoplasm (*Neiman, 2011*), we predicted that expressing extra Wtf4^antidote^ prior to spore formation from the *p^poison^* promoter could lead to greater Wtf4^poison^ exclusion from spores. To test this, we imaged protein localization in asci generated by *p^poison^_wtf4^antidote^-GFP*/ *mCherry-wtf4^poison^* diploids (Fig 3E). Inconsistent with our hypothesis, we found that the amount of Wtf4^poison^ packaged in spores was unchanged by the presence of the Wtf4^antidote^ at the time of spore formation: 59% of mCherry-Wtf4^poison^ was contained in spores in asci generated from *mCherry-wtf4^poison^/ p^poison^_wtf4^antidote^-GFP* (Fig 3D-left), while 55% was in spores in asci generated from *mCherry-wtf4^poison^/ ade6+* diploids (Fig 3D-middle). Similar to our experiments described above, we found that a significantly greater fraction of the mCherry-Wtf4^poison^ (59%) was packaged in spores relative to the Wtf4^antidote^-GFP (47%) in asci generated by *p^poison^_wtf4^antidote^-GFP*/ *mCherry-wtf4^poison^* diploids (Fig 3D-left, p< 0.05, paired t-test). We note that, in experiments discussed below, we did find that expression of Wtf4^antidote^ from its endogenous promoter was able to increase the fraction of Wtf4^poison^ protein excluded from spores (Fig 4D and 4E).

**Figure 4.**
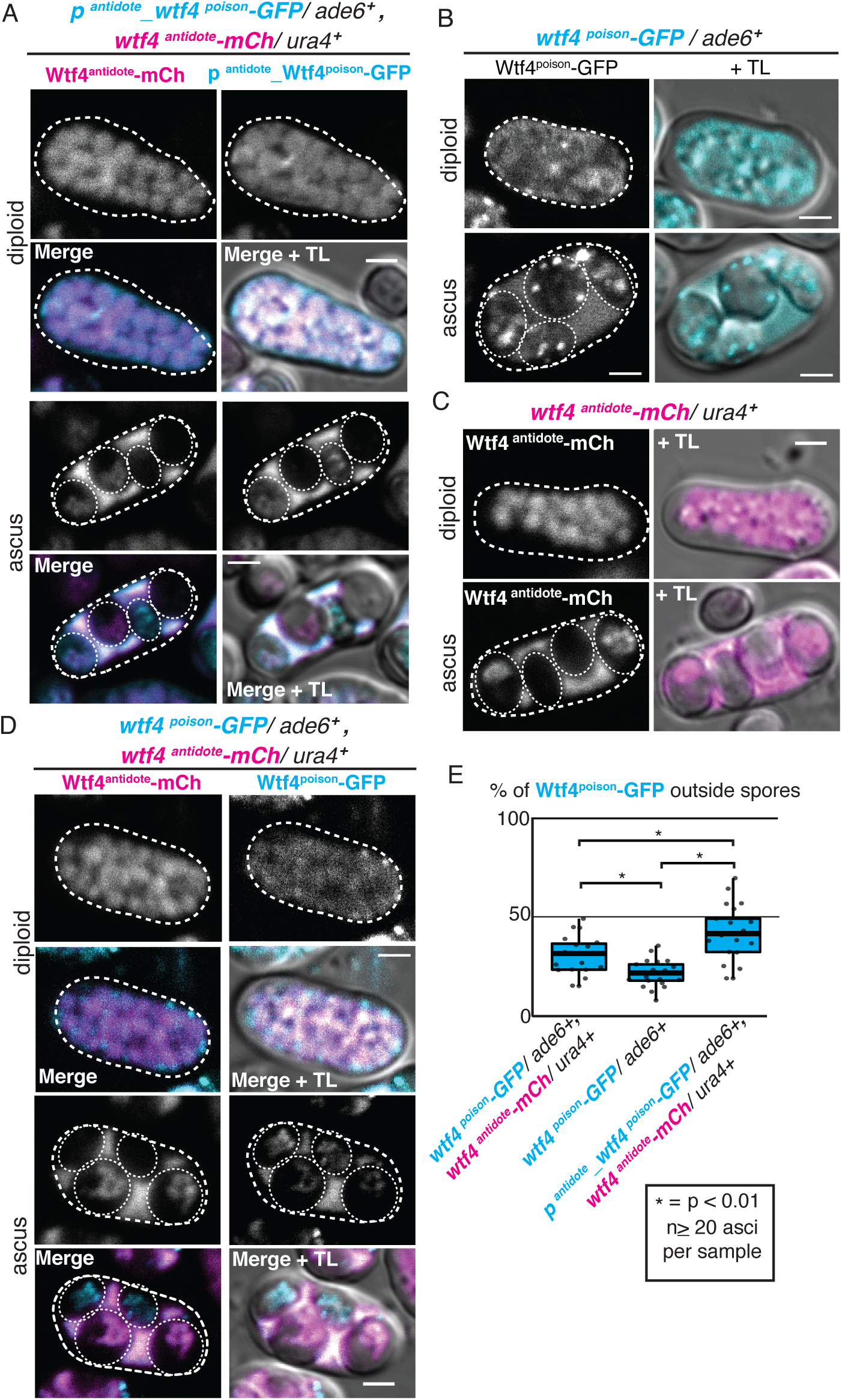
Wtf4^poison^ localization is altered when expressed from the *p^antidote^* promoter. (**A**) Image of a *p^antidote^*_*wtf4^poison^-GFP/ ade6+, wtf4^antidote^-mCherry/ura4+* diploid and tetratype ascus. p^antidote^_Wtf4^poison^-GFP is shown in cyan and Wtf4^antidote^-mCherry is shown in magenta in merged images. Additional images can be found in S4B Fig. (**B**) Image of a *wtf4^poison^-GFP/ ade6+* diploid and ascus. Images of this strain were first presented in (*Nuckolls et al., 2017*). (**C**) Image of a *wtf4^antidote^-mCherry/ ura4+* diploid and ascus. Additional images of this cross are also presented in S1C Fig. (**D**) Images of a heterozygous *wtf4^antidote^-mCherry/ura+, wtf4^poison^-GFP/ade6+* diploid and ascus. Wtf4^antidote^-mCherry is shown in magenta and Wtf4^poison^-GFP is shown in cyan in merged images. Images of this cross are also depicted in Fig 1D. (**E**) Quantification of the percent of total ascal Wtf4^poison^-GFP fluorescence located outside spores in asci generated by the following diploids: 1. *wtf4^poison^-GFP/ade6+, wtf4^antidote^-mCh/ura4+*, 2. *wtf4^poison^-GFP/ade6+,* and 3. *p^antidote^_wtf4^poison^-GFP/ade6+, wtf4^antidote^-mCh/ura4+* (*=p < 0.01, t-test, n≥ 20 asci per sample). TL= transmitted light. All scale bars represent 2 μm. All images were acquired after 3 days on sporulation media. Not all images are shown at the same brightness and contrast to avoid over saturation of pixels in the brighter images.

Given that the protein generated by the *p^poison^_wtf4^antidote^-GFP* allele was at least partially excluded from spores (Fig 3B) and was unable to fully prevent mCherry-Wtf4^poison^ from entering spores (Fig 3D and 3E), we did not expect the allele to fully protect spores from destruction by Wtf4^poison^. To test this, we assayed allele transmission in *p^poison^_wtf4^antidote^-*GFP/*wtf4* heterozygotes. If the Wtf4^antidote^ encoded by *p^poison^_wtf4^antidote^-GFP* effectively neutralizes Wtf4^poison^, the *p^poison^_wtf4^antidote^-GFP* allele should suppress drive of the wild-type *wtf4* allele (e.g. S1 Fig, diploid 13). We did not, however, observe suppression by the *p^poison^_wtf4^antidote^-GFP* allele and instead observed strong drive of *wtf4* (S4A Fig, diploid 16).

We also tested the ability of the *p^poison^_wtf4^antidote^-GFP* allele to drive in the presence of a *wtf4^poison^* allele (S4 fig, diploid 15) because a *wtf4^antidote^* allele expressed from the endogenous promoter can drive in the presence of a *wtf4^poison^* allele (e.g. S4A Fig, diploids 12 and 17, *Nuckolls et al., 2017*). In the absence of a *wtf4^antidote^* allele, we previously observed that an untagged *wtf4^poison^* allele shows drag, or is transmitted to less than half of the spores produced by a heterozygote (*Nuckolls et al., 2017*). We assume this is because spores that inherit the allele are more likely to be killed than those that do not. We also observed drag of the mCherry- *wtf4^poison^* allele (S4A Fig diploid 10). This drag was stronger in diploids also containing the *p^poison^_wtf4^antidote^-GFP* allele (i.e. *mCherry-wtf4^poison^*/*p^poison^_wtf4^antidote^-GFP*; S4A Fig, compare diploids 10 and 15). This is consistent with the *p^poison^_wtf4^antidote^-GFP* allele providing some protection to the spores that inherit it, though it is less than that provided by a *wtf4^antidote^-GFP* allele driven by the endogenous promoter (S4A Fig, compare diploids 13 and 16.)

Overall, our results suggest that transcription of *wtf4^antidote^* from its endogenous promoter in spores is required for it to fully neutralize the Wtf4^poison^. We show this is likely because a significant portion Wtf4^antidote^ produced prior to spore formation is excluded from developing spores. An alternate interpretation of our results is that the Wtf4^antidote^ protein present at the time of spore formation is not excluded from spores, but is instead degraded within spores before it can be visualized. We favor the first model, because we generally detect high levels of Wtf4^antidote^ protein in spore vacuoles (*Nuckolls et al., 2020*), so it does not seem like the turnover of the protein within spores is particularly rapid.

### Early transcription of the *wtf4^poison^* is required for efficient drive

To test if transcriptional timing of the *wtf4^poison^* was important for meiotic drive, we generated a clone, *p^antidote^_wtf4^poison^-GFP*, which swapped the *p^poison^* promoter for the *p^antidote^* promoter. However, we failed to transform this construct into cells not carrying a *wtf4^antidote^* allele, despite multiple attempts and using multiple transformation protocols (standard lithium acetate and electroporation; *Gietz, et al., 1995; Murray, Watson, and Carr, 2016*). We were also unable to lose or cross out the *wtf4^antidote^* allele in strains also carrying *p^antidote^_wtf4^poison^-GFP*. These results suggest that the *p^antidote^* promoter may cause some Wtf4^poison^ expression during the transformation process or in vegetative growth, inhibiting successful transformation of the cassette without the presence of the antidote. Consistent with this, published long-read sequencing data shows some transcription of *wtf^antidote^* alleles occurs in cells prior to meiotic induction (*Kuang et al., 2017*).

We, therefore, transformed the *p^antidote^_wtf4^poison^-GFP* cassette into the *ade6* locus of cells carrying *wtf4^antidote^-mCherry* allele at the *ura4* locus and analyzed phenotypes of cells heterozygous for both alleles. We found that Wtf4^poison^-GFP expressed from the *p^antidote^* promoter was enriched in two of the four spores (Fig 4A and S4B Fig), with similar localization to Wtf4^antidote^-mCherry (Fig 4A and 4C), while Wtf4^poison^-GFP expressed from its endogenous promoter was more evenly distributed between all four spores (Fig 4B).

In addition, when Wtf4^antidote^-mCherry and Wtf4^poison^-GFP proteins are both expressed from the *p^antidote^* promoter, we observed strong co-localization of the two Wtf4 proteins within diploid cells induced to undergo meiosis (Fig 4A). Based on our previous observations (*Nuckolls et al., 2020*) and the appearance of the cells, it is likely the colocalized proteins are within vacuoles. Our images also suggested that more of the total Wtf4^poison^-GFP was being excluded from spores in comparison to wild-type diploids where Wtf4^poison^ is under its endogenous promoter (compare Fig 1D to Fig 4A and S4B Fig). To test this hypothesis, we quantified the percent of the total ascus Wt4^poison^-GFP fluorescence signal in spores. We found that 29% of the Wt4^poison^-GFP signal was outside spores in asci generated by diploids heterozygous for both *wtf4^poison^-GFP* and *wtf4^antidote^-mCherry.* This was significantly more than the fraction of Wt4^poison^-GFP outside spores in the absence of Wtf4 antidote (21%, Fig 4E). Moreover, when the Wt4^poison^-GFP was expressed from the *p^antidote^* promoter, we observed an even larger fraction (41%) of the Wt4^poison^- GFP signal outside spores (Fig 4E). These results suggests that Wtf4^antidote^ expressed from its endogenous promoter helps exclude Wtf4^poison^ from spores, particularly if the Wtf4^poison^ is expressed from the same *p^antidote^* promoter.

We also assayed allele transmission from cells heterozygous for both *p^antidote^_wtf4^poison^-GFP* and *wtf4^antidote^-mCherry.* We found that drive of the *wtf4^antidote^-mCherry* allele in diploids carrying the *p^antidote^_wtf4^poison^-GFP* allele was weaker than in the similar diploids in which the *wtf4^poison^-GFP* allele had its endogenous promoter (S4A Fig, diploids 17 and 18). This suggests that Wtf4^poison^- GFP expressed by the antidote promoter fails to sufficiently poison all spores. In support of that interpretation, we saw significantly more wild-type (*ura+, ade6+*) spores survived in the *p^antidote^_wtf4^poison^-GFP*/*ade6*+, *wtf4^antidote^-mCherry*/*ura4+* cross than in the *wtf4^poison^-GFP*/*ade6*+, *wtf4^antidote^-mCherry*/*ura4+* cross (28% in comparison to 5%, S4A Fig, diploids 17 and 18). This inefficient killing by Wtf4^poison^-GFP expressed under the *p^antidote^* promoter is consistent with our observations that less of the Wtf4^poison^-GFP protein enters spores, relative to the protein expressed from its endogenous promoter. Altogether, our results show that transcription of the *wtf4^poison^* from its endogenous promoter is required for optimal function in meiotic drive.

### Wtf4^antidote^ promoter is induced by ER stress

As mentioned above, we failed to transform the *p^antidote^_wtf4^poison^-GFP* construct into cells not carrying a *wtf4^antidote^* allele, but we could introduce it to strains carrying *wtf4^antidote^-mCherry.* We hypothesized that the stress of transformation could be increasing the expression of the Wtf4^poison^ from the *p^antidote^* promoter, leading to cell death. Consistent with this idea, it was previously observed that the *wtf4* allele from the common lab isolate of *S. pombe* has slightly increased expression under heat stress and hydrogen peroxide treatment (*Chen et al., 2003*).

To formally test if the *p^antidote^* promoter from *S. kambucha wtf4* was upregulated under different stress conditions, we employed a transcriptional reporter. Specifically, we integrated the *p^antidote^_mCherry* reporter allele described above at the *ura4* locus. We then compared the fluorescence of mCherry from cells exposed to varying growth conditions. We found that expression of the *p^antidote^_mCherry* was not significantly increased after 1 hour of heat shock (40°C) relative to growth at the standard temperature (32°C). We did, however, observe a significant increase in *p^antidote^_mCherry* expression in cells after 1 hour of growth in 0.5 µg/ml tunicamycin, which induces endoplasmic reticulum stress, relative to untreated cells (S5 Fig, p< 0.05, t-test). These results support the idea that the *p^antidote^* promoter can be expressed outside of sporulation and upregulated in response to stress.

### Master meiotic regulator, Mei4, controls Wtf4^poison^ expression

Transcriptional regulation of the *p^poison^* promoter is particularly important because it induces the production of a toxic protein. We reasoned that if cells could prevent transcription from *p^poison^* promoters, they could suppress drive and increase fertility of cells heterozygous for *wtf* drivers. Previous work noted that the transcription of multiple *wtf* genes was controlled by the fork-head transcription factor Mei4 as *wtf* transcription is decreased if *mei4* is deleted and *wtf* gene expression is increased if Mei4 is overexpressed (*Mata et al., 2007*). Mei4 controls the expression of hundreds of genes and is known as the master regulator of middle meiosis genes (*Ioannoni, et al., 2016; Alves-Rodrigues et al., 2016*). However, the prior study was performed before the discovery that *wtf* drivers can make two transcripts, so it was not clear which transcript Mei4 controls.

To understand which transcript Mei4 controls, we first looked for the Mei4-binding motif in the *wtf4* promoters. Fork-head transcription factors bind sequences containing FLEX motifs and the complete Mei4 binding motif contains a nine-base pair (GTAAACAAA) core sequence (*Alves-Rodrigues et al., 2016; Horie et al., 1998*; *Ioannoni, et al., 2016; Moldón et al., 2008*). We found this nine-base pair Mei4 binding motif in the *p^poison^* promoter 110 base pairs upstream of the *wtf4^poison^* transcriptional start site, in reverse orientation to the TSS (Fig 5A). Moreover, we found this sequence was conserved amongst known *wtf* drivers (Fig 5A).

**Figure 5.**
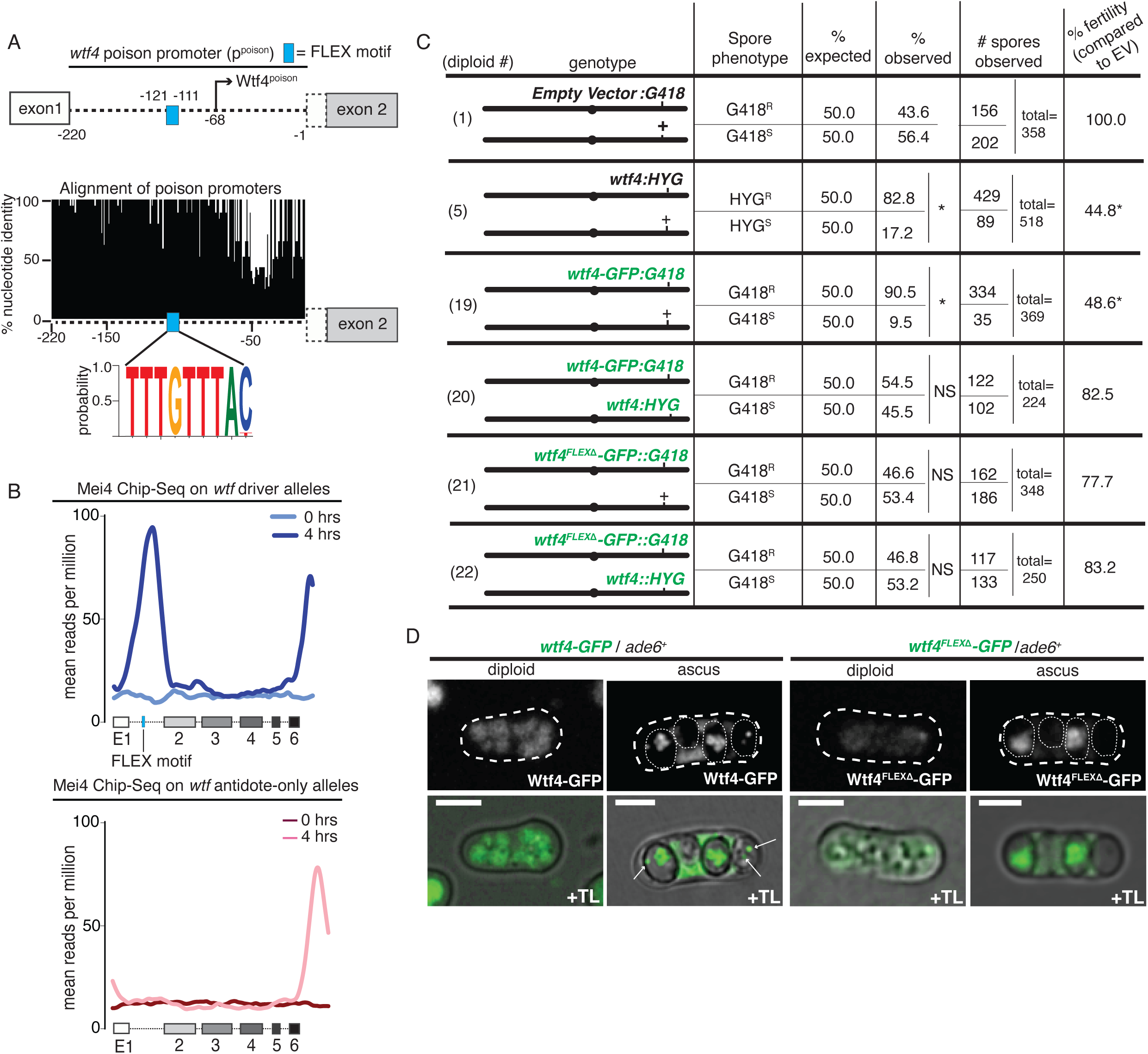
Master meiotic regulator, Mei4, controls Wtf4^poison^ transcription. (**A**) Depiction of the *wtf4*^poison^ promoter contained within intron 1. There is a FLEX motif (blue box, TGTTTAC in opposite orientation) located 111 to 117 nucleotides upstream of the Wtf4^poison^ transcriptional start site. Percent consensus nucleotide identity of intron 1 sequences of 28 predicted poison-antidote *wtf* drivers (*Eickbush et al., 2019*) from three different strains of *S. pombe* (the reference genome*, S. kambucha,* and *FY29033*). The figure shows the percent consensus identity at each nucleotide position, excluding gaps. Predicted pseudo-genes and the highly diverged *wtf* genes of unknown function were excluded (*Eickbush et al., 2019*). (**B**) Mei4 Chip-seq data (data from *Alves-Rodrigues et al. 2016*) showing Mei4 binding on *wtf* genes at both 0 hours and 4 hours after meiotic induction. On the top panel, the reads are aligned to the predicted *wtf* meiotic drive genes (*wtf4, wtf13*) and on the bottom the reads are aligned to the *wtf* genes that are predicted to only encode antidote proteins (*wtf5, wtf9, wtf10, wtf16, wtf18, wtf20, wtf21, wtf25*) in the *S. pombe* genome version ASM294v2. For any reads that mapped to more than one location, only a single location, chosen at random, is reported. (**C**) Allele transmission and fertility (assayed via <u>v</u>iable <u>s</u>pore <u>y</u>ield, VSY) of diploids with the depicted genotype. Alleles listed are at *ade6.* The genotype column shows a cartoon depiction of the relevant genotype. The progeny phenotypes are then shown on the right. For diploids heterozygous at one locus (e.g. Diploid 1), two values are shown (top and bottom) that represent the two possible haploid genotypes. Spores exhibiting both parental phenotypes were considered diploid or aneuploid and were excluded from this table but can be found in S4 Data. The expected values assume Mendelian allele transmission. (*= p < 0.05, NS= not significant; G-test for allele transmission, Wilcoxon test for VSY, in comparison to the empty vector control). We compared all diploids to diploid 1 as the control. The data for diploid 1 is also depicted in S1A Fig and S4A Fig. Diploid 5 is also shown S1A Fig. (**D**) Images of heterozygous *wtf4-GFP/ade6+* (left) and *wtf4^FLEX^*^Δ^*-GFP/ade6+* (right) diploids and asci acquired after 2 days on sporulation media. Wtf4-GFP and Wtf4^FLEXΔ^ -GFP are shown in green. Scale bar represents 4 µM. TL= transmitted light. Not all images are shown at the same brightness and contrast to avoid over saturation of pixels in the brighter images.

To test if Mei4 binds this motif in meiotic cells, we analyzed data from a previous Chromatin-Immunoprecipitation sequencing (ChIP-seq) experiment of Mei4 during meiosis done in the lab isolate of *S. pombe* (*Alves-Rodrigues et al., 2016*). Due to the similarity of the *wtf* genes, many reads could not be uniquely assigned to *wtf4* or any single *wtf* gene and were thus randomly assigned to matching sites. The sequence of intron 1, however, is distinct between *wtf* drivers that contain the *p^poison^* promoter and antidote-only *wtf* genes that do not contain the *p^poison^* promoter, allowing us to distinguish Mei4 binding between the two classes of *wtf* genes (*Bravo Núñez et al., 2018b; Eickbush et al, 2019, Hu et al., 2017*). We observed that Mei4 binding to *wtf* drivers (*wtf4*, *wtf13*) was low, relative to the genome average, prior to meiosis (Fig 5B, 0 hours). After 4 hours in meiosis, when most cells are in prophase I, we saw a strong increase in reads in the *p^poison^* promoter (intron 1) of the *wtf* drivers (Fig 5B, 4 hours). We did not see an increase in Mei4 binding to antidote-only *wtf* genes (*wtf5*, *wtf9*, *wtf10*, *wtf16*, *wtf18*, *wtf20*, *wtf21*, and *wtf25*) during meiosis (Fig 5B). We also saw a Mei4 peak at the C-terminal region of both subsets of *wtf* genes. We predict this peak is due to the nearby LTRs, as we saw high Mei4 binding on LTRs independently of their association with *wtfs* (data not shown). Together, these results suggest that Mei4 binds to the *p^poison^* promoters of *wtf* drivers to transcribe *wtf4^poison^*.

To test this idea further, we deleted the FLEX motif from the *wtf4-GFP* allele described above to generate *wtf4^FLEX^*^Δ^*-GFP*. We then integrated this allele at *ade6* to test its function. We did not anticipate that deletion of the Mei4-binding motif would affect the expression or function of the Wtf4^antidote^ protein as we can delete all of intron 1 from *wtf4* (as in *wtf4^antidote^-mCherry*), without affecting Wtf4^antidote^ function (S1A Fig, diploids 6 and 7). Consistent with our expectations, the *wtf4^FLEX^*^Δ^*-GFP* allele encodes a functional antidote as it suppresses drive of the wild-type gene in a *wtf4^FLEX^*^Δ^*-GFP* /*wtf4+* heterozygote (Fig 5C, diploid 22). We tested the ability of the *wtf4^FLEX^*^Δ^*-GFP* allele to encode a functional Wtf4^poison^ by assaying if the allele could drive in a w*tf4^FLEX^*^Δ^-GFP / *ade6+* heterozygote. These diploids had wild-type fertility and Mendelian allele transmission (Fig 5C, diploid 21). As the w*tf4^FLEX^*^Δ^-GFP does encode a functional antidote, we conclude that the allele does not drive because the Mei4-binding motif is essential for production of the Wtf4^poison^. This was supported by the fact that when we imaged *wtf4^FLEX^*^Δ^- GFP/*ade6+* diploids, we saw strong GFP signal within two of the four spores but did not observe the irregularly sized puncta that are characteristic of the Wtf4^poison^ protein. In contrast, the asci generated from *wtf4-GFP*/*ade6+* had both the foci within the spores and puncta throughout (Fig 5D, arrows point to puncta typical of Wtf4^poison^). These data support that Mei4 controls expression of the Wtf4^poison^ and likely the poison proteins produced by other *wtf* drivers.

## Discussion

### Dual promoters with distinct regulation are key to *wtf* meiotic drive

In order for *wtf4* to drive, all spores must be exposed to a lethal dose of the Wtf4^poison^, while only those that inherit the *wtf4* allele can receive a sufficient dose of the Wtf4^antidote^ to survive. Previously, we observed Wtf4^poison^ in meiotic cells prior to spore formation, but observed Wtf4^antidote^ enriched only in spores (*Nuckolls et al., 2017*). These observations explained how all spores were poisoned and how only those that inherited the *wtf4* locus were rescued by the antidote, but it raised a new puzzle of how meiotic cells survived exposure to the Wtf4^poison^ in the apparent absence of the Wtf4^antidote^. This uncertainty was amplified by subsequent work that showed Wtf4^poison^ efficiently kills cells, not just spores, that do not express antidote (*Nuckolls et al., 2020*). In this work, our new data offer a solution to the puzzle in that we found there is also Wtf4^antidote^ expression in cells undergoing meiosis. This result is consistent with previous long-read RNA sequencing data that show a low level of transcription of *wtf^antidote^* alleles in early meiosis (*Kuang et al., 2017*). The Wtf4^antidote^ protein we observe in cells undergoing meiosis may be produced using the second viable start codon in the Wtf4^antidote^ coding sequence, because we observe the Wtf4^antidote^ in diploids with C-terminal fluorescent tags, but not with an N-terminal tag in front of the first start codon (S1B-D Fig).

Our work also reveals that different transcriptional regulation of the poison and antidote promoters is largely responsible for the distribution of the Wtf4^poison^ to all spores and the enrichment of the Wtf4^antidote^ in the spores that inherit the locus, as transcriptional reporters grossly recapitulate localization patterns of tagged proteins (Fig 2). However, different features of the proteins also contribute to their localization patterns. Specifically, we found that the propensity for the two Wtf proteins to be included in developing spores was different. The Wtf4^antidote^ present in the cytoplasm at the time of spore encapsulation is largely excluded from the spores. In the absence of the Wtf4^antidote^, the Wtf4^poison^ is included in the developing spores (Fig 4E). In the presence of the Wtf4^antidote^ expressed from its endogenous promoter, we observed that the amount of Wtf4^poison^ included in developing spores was decreased, but not eliminated (Fig 4E).

When both the Wtf4^poison^ and Wtf4^antidote^ proteins were expressed from the same promoters, our results were not entirely consistent. When the two proteins were expressed from the *p^antidot^*^e^ promoter, we found that the Wtf4^poison^ and Wtf4^antidote^ colocalized (likely in vacuoles) and that a larger fraction of the of the Wtf4^poison^ was excluded from spores relative to when only Wtf4^poison^ is present at the time of spore formation (Fig 4E). This result is consistent with our previous observations of Wtf4 proteins in vegetatively growing *S. pombe* or *S. cerevisiae* cells where the two proteins coassemble and are trafficked to the vacuole (*Nuckolls et al., 2020*). When the Wtf4^poison^ and Wtf4^antidote^ were both expressed from the p^poison^ promoter, however, we did not observe a larger fraction Wtf4^poison^ excluded from the spores relative to when only the Wtf4^poison^ was present (Fig 3D). The reasons for this discrepancy are unclear. We speculate that there could be additional factors (e.g. potential cofactors or precise expression level) that promote the inclusion of the Wtf4^poison^ in spores and promote the ability of the Wtf4^antidote^ to sequester Wtf4^poison^ outside of spores that work best when the proteins are expressed by their endogenous promoters.

We show that transcriptional timing and selective protein inclusion in spores are essential for the proteins to coordinate drive. Our data suggest the following model (Fig 6): the Wtf4^poison^ is mostly expressed prior to spore formation and then subsequently packaged in spores. This ensures that each spore gets a lethal dose from a heterozygous zygote. Expression of the poison from the antidote promoter is insufficient to poison all spores (S4A Fig, diploid 18). Conversely, Wtf4^antidote^ is present prior to spore formation, perhaps just enough to prevent the cells undergoing meiosis from succumbing to the poison (S1C-D Fig). The antidote that is present at the time of spore formation likely offers little protection to spores that do not inherit the driving locus because it does not prevent all Wtf4^poison^ from entering spores (Figure 4E) and the Wtf4^antidote^ is excluded from developing spores (Fig 3A-B). Because of these factors, spores that inherit *wtf4* must produce their own private supply of Wtf4^antidote^, while those that do not succumb to the poison.

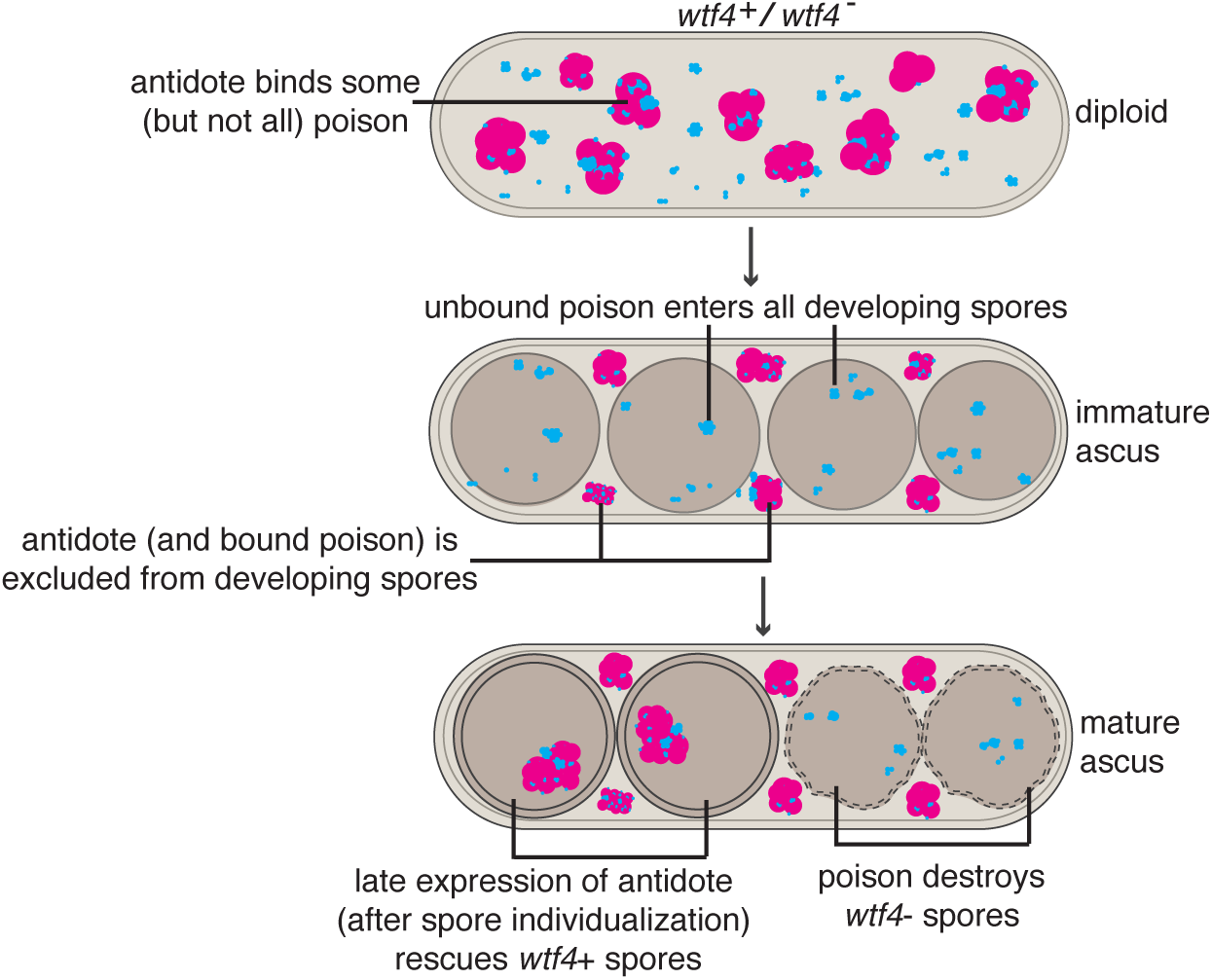

### Difficulties in universal suppression of *wtf* drivers

This work also adds to our understanding of the evolutionary success of the *wtf* drivers within *S. pombe*. All sequenced isolates of *S. pombe* contain between 25 and 38 *wtf* genes, including 4-14 genes predicted or shown to be intact meiotic drivers (*Bravo Núñez et al., 2020a*; *Eickbush et al., 2019, Hu et al., 2017; Nuckolls et al 2017*). Each heterozygous driver can cause the destruction of half the spores and antidote proteins generally do not protect against poisons that have distinct sequences (*Bravo Nunez et al., 2020, Hu et al., 2017*). Because of these features, diploid zygotes produced by crossing two diverged isolates produce very few viable spores (*Bravo Núñez et al., 2020a*; *Gutz and Doe 1975*; *Hu et al., 2017; Jeffares et al., 2017; Nuckolls et al., 2017; Singh and Klar, 2002; Tau et al., 2019; Zanders et al., 2014*). The incentive for *S. pombe* to evolve a suppression mechanism to suppress a *wtf* driver, therefore seems quite high (*Crow, 1991*).

There are likely many roads to suppression of *wtf* drivers, but we speculate that none of them have been terribly successful given the prevalence and high activity of these genes in *S. pombe*. The known genic suppressors of *wtf* drivers are all Wtf^antidote^ proteins. Importantly, Wtf^antidote^ proteins seem highly specific in that they neutralize only Wtf^poison^ proteins with very similar sequences (*Hu et al., 2017*, *Bravo Núñez et al., 2018a*, *Bravo Núñez et al., 2020a*, *Nuckolls et al., 2020*). Because of this, Wtf^antidote^ proteins are not expected to act as general suppressors of *wtf* drivers, as the genome would require a unique suppressor for each driver. A non-Wtf universal suppressor that interfaces with Wtf^poison^ proteins might also be difficult to envision given the vast diversity of Wtf proteins, which can share as little as 30% amino acid identity (*Bravo Núñez et al., 2020a, Eickbush et al., 2019*).

Transcriptional repression seems like a more feasible route to universal suppression of *wtf* drivers given the conservation of their promoters (*Bowen et al., 2003; Eickbush et al., 2019; Hu et al., 2017;* Fig 2B and 2D). Previous work has identified some factors that help regulate the expression of *wtf* genes, however, these studies did not distinguish between *wtf^poison^* and *wtf^antidote^* transcripts (*Hansen et al., 2005; Watts et al., 2018*). In mitotic cells, a mutation in the histone deacetylase *clr6* or inhibition of histone deacetylation with the drug Trichostatin A leads to increased *wtf* gene transcription (*Hansen et al., 2005; Nicolas et al., 2007; Watts et al., 2018, Zilio et al., 2014*). Also in mitotic cells, *wtf* mRNAs can be eliminated from cells via double-stranded-directed RNA decay (*Sugiyama and Sugioka-Sugiyama, 2011*). It is possible that these mitotic mechanisms may affect *wtf* gene expression during meiosis as well, but this has not been determined. In meiotic cells, Cuf2 decreases the expression of *wtf* genes late in meiosis (*Ioannoni et al., 2012, Ioannoni et al., 2016*). However, this transcriptional suppression appears to have a minimal effect on drive. For example, Cuf2 decreases expression of *wtf13*, but even in this suppressed state, *wtf13* drives into >90% of the gametes produced by a *wtf13+/wtf13*Δ heterozygote (*Bravo Núñez et al., 2018a*).

Effective transcriptional repression of *wtf* drivers could be challenging for several reasons. The first challenge is the large number of *wtf* genes and the fact that they are not all found at the same locus. This may make it hard to simply turn them all off *en masse* by packaging one genomic region in heterochromatin. Even if the genes were found in one location, the genomic region housing the *wtf* genes would likely be under strong selection to resist such heterochromatization. Specifically, haplotypes that could avoid heterochromatization and thus better maintain drive would be selected by drive. This is because regions linked to meiotic drivers also get to enjoy the benefits of drive and thus profit from enhancing drive, rather than suppressing it (*Crow, 1991*). This selective pressure resisting heterochromatization would also apply to the distributed *wtf* drivers as well. This situation is profoundly different than loci housing transposable elements as those selfish elements generally offer no evolutionary advantages to flanking sequences (*Burt and Trivers, 2006*).

Another challenge in establishing suppression of *wtf* drivers is that partial or imperfect suppression of *wtf* genes may do more harm than good. For example, if silencing is incomplete in a diploid homozygous for a given driver, only one allele could be expressed, and drive would occur in a cell where no drive would occur in the absence of silencing. In addition, in some instances *wtf* drivers are suppressed by other *wtf* genes (*Bravo Núñez et al., 2018a, Bravo Núñez et al., 2020a*). This is an additional situation in which incomplete silencing of the wrong *wtf* gene could lead to more, rather than less, drive.

An ideal suppression approach would be to suppress a key transcription factor driving expression from *p^poison^* promoters. This approach would allow the cell to stop drive, but it also comes with minimal risk as partial suppression would still be selectively advantageous by decreasing the likelihood of drive. In this scenario, an imperfect system could adapt and improve over time. In this work, we identified Mei4 as the key transcription factor governing the expression of *p^poison^* (Fig 5). Suppression of Mei4, however, is not an ideal option for suppressing drive because it is an essential regulator of meiosis. Mei4 controls the expression of over 100 genes and cells lacking Mei4 fail to complete meiosis (*Abe and Shimoda, 2000; Alves-Rodrigues et al., 2016; Horie et al., 1998; Ioannoni, et al., 2016; Murakami-Tonami et al., 2007; Shimoda et al., 1985; Ruan et al., 2015; Moldón et al., 2008*).

### Final speculation

It is hard not to admire how beautifully adapted the *wtf* parasites are at promoting themselves despite the costs to *S. pombe*. Equally remarkable is how well insulated *wtf* parasites appear to be against *S. pombe* acting to stop their expression. That is not to say, however, that *S. pombe* has not found other ways to mitigate the costs of *wtf* drivers. Some *S. pombe* natural isolates preferentially inbreed and some isolates generate disomic (aneuploid or diploid) spores at high frequencies (up to 46% of spores; *López Hernández et al., 2021, Bravo Núñez et al., 2020b*). Although it is hard to say if the *wtf* genes promoted the evolution of these traits, it is possible, as both traits effectively suppress fitness costs of *wtf* drivers (*López Hernández et al., 2021*, *Bravo Núñez et al., 2020b*). If these traits were selected for based on their ability to suppress drive, suppression likely came at a steep price to fitness, given that inbreeding and meiotic disruption are generally not good for fitness. Perhaps this was the price that had to be paid, as cheaper options, like universal transcriptional silencing, were out of reach.

## Materials and methods

### Generation of yeast strains

We confirmed the plasmids generated in this section via Sanger sequencing. We completed all transformations described in this section using standard lithium acetate protocol (*Gietz, et al., 1995*), first selecting for drug resistance and then screening for the relevant auxotrophy.

#### Generation of an allele of wtf4 with endogenous promoters and mCherry-wtf4^poison^

We ordered a gBlock from Integrated DNA technologies (IDT, Coralville IA) containing the *p*^antidote^ promoter (- 600 bp), exon 1, intron 1, and sequence encoding an mCherry tag and a 5X glycine linker upstream of the *wtf4^poison^* start codon (*Hailey et al., 2002*). The gBlock also included 177 bp of exon 2. We amplified this gBlock using oligos 620 and 718. We also amplified the rest of the *wtf4* sequence from pSZB189 using oligos 679 and 687 (*Nuckolls et al., 2017*). We then used overlap PCR to stitch the PCR fragments together using oligos 620 and 687. Next, we digested the resulting PCR product with SacI and ligated it into the SacI site of pSZB188 (*Nuckolls et al., 2017*), an *ade6* integrating vector with a *kanMX4* cassette, to create pSZB250. We cut pSZB250 with KpnI and transformed the linearized vector into SZY643 to generate SZY1037.

#### Generation of a wtf4 allele that encodes only mCherry-wtf4^poison^ under the control of the endogenous promoter

This allele has the full-length *wtf4* gene (including *p^antidote^* promoter) but the two start codons in exon 1 (for the Wtf4^antidote^) are mutated to TAG. We first cloned pSZB259 which contains an untagged *wtf4* allele in which both start codons are mutated, similar to pSZB257 (*Nuckolls et al., 2017*). We amplified the *p^antidote^* promoter and the mutated version of exon 1 from pSZB259 using oligos 620 and 861. We amplified the *mCherry-wtf4^poison^* sequence from pSZB250 (described above) using oligos 862 and 687. We then used overlap PCR to stitch the two PCR products together using oligos 620 and 687. We digested the resulting PCR product with SacI and ligated the cassette into the SacI site of pSZB188 (*Nuckolls et al., 2017*) to make pSZB355. We also digested pSZB355 with SacI to isolate the *wtf4 allele* and cloned it into SacI-digested pSZB386, a hygromycin B-resistant *ade6* integrating vector (*Nuckolls et al., 2017*) to generate pSZB824. We digested pSZB824 with KpnI to linearize the construct and transformed into SZY2572 to generate SZY4570.

#### Generation of a wtf4 allele that encodes only wtf4^antidote^-GFP under the control of the endogenous promoter

We used pSZB203 as a template to amplify the 5’ end of *wtf4-GFP* with oligos 620 and 736 and the rest of the gene with oligos 735 and 634 (*Nuckolls et al., 2017*). The two PCR products were then stitched together using overlap PCR using oligos 620 and 634. The 735 and 736 oligos mutated the start codon for the *wtf4^poison^* to TAC. The resulting PCR product was then digested with SacI and ligated into the SacI site of pSZB188 to generate pSZB260. We cut pSZB260 with KpnI and integrated it into the *ade6* locus of SZY643 to create SZY1056.

#### Generation of a wtf4 allele that encodes only mCherry-wtf4^antidote^ under the control of the endogenous promoter

We used site-directed mutagenesis to mutate the translational start site from ATG to TAC of *wtf4^poison^* within pSZB248 (*Nuckolls et al., 2017*) to generate pSZB367. We linearized pSZB367 with KpnI and integrated it into SZY643 to make SZY3586.

#### Generation of a p^antidote^_mCherry transcriptional reporter

We first amplified the *wtf4^antidote^* promoter linked to mCherry using pSZB248 (*Nuckolls et al., 2017*) as a template with oligos 688 and 1447. Next, we amplified the *ADH1* transcriptional terminator from pKT127 (*Sheff et al., 2004*) using oligos 1448 and 634. We then used overlap PCR to unite these fragments and generate *p^antidote^_mCherry*. We then digested the complete *p^antidote^_mCherry* cassette with SacI and ligated it into the SacI site of pSZB386 (*Bravo Núñez et al., 2018a*) to create pSZB766. We cut pSZB766 with KpnI and integrated it into the *ade6* locus of SZY2080 to create SZY2137. We also digested the cassette form pSZB766 and cloned it into SacI-digested pSZB331 (*Bravo Núñez et al., 2020a*) to create pSZB744. We cut pSZB744 with KpnI and integrated it into the *ura4* locus of SZY2080 to create SZY4534.

#### Generation of a p^antidote long^_mCherry transcriptional reporter

We amplified the majority of the *p^antidote long^_mCherry* construct (with an *ADH1* transcriptional terminator) using oligos 3024+634 and pSZB891 (*Nuckolls et al., 2020*) as a template. We then added the rest of the upstream to the *p^antidote long^_mCherry* construct by using it as a template for PCR with oligos 3025+634. This generated the full *p^antidote long^-mCherry* construct. We digested this cassette with SacI and ligated it into the SacI site of pSZB322 (*Bravo Núñez et al., 2018*) to create pSZB1361. We cut pSZB1361 with KpnI and integrated into the *lys4* locus of SZY2080 (*Nuckolls et al., 2020*) to generate SZY4442.

#### Generation of a strain with wtf4::hphMX6 at the ade6 locus

We digested pAG32 (*Goldstein and McCuster, 1999*) with NotI to isolate the *hphMX6* cassette. We transformed the *hphMX6* cassette into SZY887 (*Nuckolls et al., 2017*) selecting for HYG resistance and lack of G418 resistance. This created SZY969.

#### Generation of a p^poison^_exon1-GFP allele

We amplified from pSZB203 the *p^poison^* promoter using oligos 1174 and 1176, *wtf4* exon 1 using oligos 1175 and 1136, and GFP (with an *ADH1* transcriptional terminator) using oligos 1137 and 634 (*Nuckolls et al., 2017*). We used overlap PCR to join the three pieces using oligos 1174 and 634. We then digested the complete *p^poison^_exon1-GFP* construct with SacI and cloned it into SacI-digested pSZB386 to generate pSZB553.

#### Generation of a p^poison^_GFP transcriptional reporter

We used pSZB203 as a template to amplify the p^poison^ promoter using oligos 1174 and 1549, and to amplify GFP (with an *ADH1* transcriptional terminator) using oligos 1548 and 634 (*Nuckolls et al., 2017*). We used overlap PCR to join these two pieces using oligos 1174 and 634. We then digested the complete *p^poison^_GFP* construct with SacI and cloned it into SacI-digested pSZB386 to generate pSZB821. We linearized pSZB821 with KpnI and integrated it into SZY643 to make SZY2279.

#### Generation of an allele of wtf4^antidote^-GFP in which the p^antidote^ promoter is replaced with the p^poison^ promoter

We amplified the poison promoter + exon 1 from pSZB553 (see above) using oligos 1174 and 604. We amplified the rest of the *wtf4* coding sequence from pSZB700 (*Nuckolls et al., 2020*) using oligos 605 and 997. We amplified GFP (with an *ADH1* transcriptional terminator) from pSZB203 (*Nuckolls et al., 2017*) using oligos 998 and 634. We used overlap PCR to join the three pieces using oligos 1174 and 634.We digested the complete *p^poison^_wtf4^antidote^-GFP* construct with SacI and cloned it into SacI-digested pSZB386 to generate pSZB727. We digested pSZB727 with KpnI to linearize the construct and transformed into SZY44 to generate SZY2406.

#### Generation of the p^antidote^_wtf4^poison^-GFP allele

We amplified *p^antidote^* from pSZB203 using oligos 688 and 1383 (*Nuckolls et al., 2017*). We amplified the *wtf4^poison^* coding sequence from pSZB392 (*Nuckolls et al., 2020*) using oligos 1384 and 997. We amplified GFP (with an *ADH1* transcriptional terminator) from pSZB203 (*Nuckolls et al., 2017*) using oligos 998 and 634. We used overlap PCR to join the three pieces using oligos 688 and 634. We digested the complete *p^antidote^_wtf4^poison^-GFP* construct with SacI and cloned it into SacI-digested pSZB386 to generate pSZB758. We digested pSZB758 with KpnI to linearize the construct and transformed into SZY2572 to generate SZY4525. The *p^antidote^_wtf4^poison^-GFP* allele lacks exon 1 and thus does not encode Wtf4^antidote^.

#### Generation the wtf4^FLEXΔ^-GFP allele

We used site-directed mutagenesis to delete the FLEX motif (TTTGTTTAC, *Horie et al., 1998*; *Moldón et al., 2008*) within intron 1 of the *wtf4-GFP* allele in pSZB203 (*Nuckolls et al., 2017*). This generated pSZB848. We digested pSZB848 with KpnI to linearize the construct and transformed into SZY643 to generate SZY1479.

#### Generation of an ade6+::his5Δ allele

This was completed in the same manner as in *Nuckolls et al., 2017*. Briefly, we amplified from genomic DNA a region upstream of *his5* using oligos 795 and 796, a region downstream of *his5* using oligos 797 and 798, and *ade6+* using oligos 799 and 800. We stitched these pieces together using overlap PCR and oligos 795 and 798. We then transformed the cassette into SZY631 (*Bravo Núñez et al., 2018a*), selecting for *ade6+* and then screening for *his5-.* This generated SZY1285 used in this work.

#### Generation of a ura4+, wtf4^poison^-GFP strain

We amplified a *ura4+* cassette using oligos 34+37 and SZY44 (*Nuckolls et al., 2017*) as a template. We then transformed it into SZY1049 (*Nuckolls et al., 2017*), selecting for *ura4+,* to generate SZY5047.

### Allele transmission and viable spore yield

All allele transmission and viable spore yield (VSY) assays were completed as previously described (*Nuckolls et al., 2017*; *Smith, 2009*). Briefly, we generated stable diploids by mixing each haploid parent in a microcentrifuge tube and plating them on SPA (1% glucose, 7.3 mM KH_2_PO_4_, vitamins, agar) for ∼15 h at room temperature to allow the cells to mate. We scraped the mated cells off of SPA and spread on a medium to select for heterozygous diploids (minimal yeast nitrogen base plates). We grew diploid colonies overnight in 5 mL of rich YEL broth (0.5% yeast extract, 3% glucose, 250 mg/L of adenine, lysine, histidine, and uracil). We then plated ∼100 uL onto SPA to induce sporulation, as well as diluted samples onto YEA (same as YEL, but with agar). We confirmed the colonies that grew on the YEA plate were truly heterozygous diploid cells by replicating to diagnostic media and counted the colonies for VSY. After 3 days, we scraped the cells from the SPA plates, treated it with glusulase (Sigma (G7017-10ML) and ethanol to isolate spores, and plated dilutions of the spores on YEA. We then counted and phenotyped the spore colonies using standard approaches. At least 2 diploids were assayed per cross and at least 150 spores were genotyped for allele transmission assays.

### Alignment of promoters

For the alignment of antidote promoters (Fig 2B), we aligned the 600 base-pairs upstream of 41 predicted antidote-only alleles (*Eickbush et al., 2019*) from three different strains of *S. pombe* (the reference genome*, S. kambucha,* and *FY29033, Lock et al., 2018*). For the alignment of poison promoters (Fig 2D), we aligned the intron 1 sequences of 28 predicted poison-antidote *wtf* drivers (*Eickbush et al., 2019*) from three different strains of *S. pombe* (the reference genome*, S. kambucha,* and *FY29033*). We utilized Geneious (version 10.0.7, https://www.geneious.com) using the Geneious aligner with the “global alignment without free end gaps” setting, taking the percent identity at each nucleotide position.

### Fluorescence microscopy

We generated diploids as previously described (*Nuckolls et al., 2017*) and placed them on sporulation agar (SPA, 1% glucose, 7.3 mM KH_2_PO_4_, vitamins, agar) for 2-3 days. We scraped the cells off of the plates and onto slides for imaging. For all microscopy, except for the experiments listed below, we used an LSM-780 (Zeiss) microscope, with a 40x C-Apochromat water-immersion objective (NA 1.2), in photon-counting channel mode with 488 and 561 nm excitation. We collected GFP fluorescence through a 481–552 bandpass filter and mCherry through a 572 long-pass filter. We also used photon-counting lambda mode, with 488 and 561 nm excitation, collecting fluorescence emission over the entire visible range. We then used these images to linearly unmix the fluorescence spectra using an in-house custom written plugin for ImageJ (https://imagej.nih.gov/ij/) to verify that there was no auto-fluorescence in the cells. Brightness and contrast are not the same for all images. We utilized two independent progenitor diploids and assayed at least 25 asci for each genotype represented. We called an ascus “mature” if all four spores showed distinct, dark outlines via transmitted light, suggesting spore membranes had been formed. For images where fluorescence intensity inside of spores is compared to fluorescence intensity outside of spores, great care was taken in acquiring the data so that only asci where all spores were sharply in focus were used for comparison.

For the *wtf4^antidote^-mCherry/ura4+* (S1C Fig) we imaged using a Zeiss Observer.Z1 wide-field microscope with a 40x C-Apochromat (1.2 NA) water-immersion objective and collected the emission onto a Hamamatsu ORCA Flash 4.0 using μManager software. We used BP 440-470 nm to excite GFP and collected BP 525-550 emission using a FT 495 dichroic, and mCherry with BP 530–585 nm excitation and LP 615 emission, using an FT 600 dichroic filter, and mCherry with BP 530–585 nm excitation and LP 615 emission, using an FT 600 dichroic filter. Brightness and contrast are not the same for all images.

For the gametogenesis time-lapse imaging (Fig 1C), we crossed a haploid *Sp* strain carrying *mCherry-wtf4* (SZY1142) to one with *wtf4^poison^-GFP* (SZY1049) to generate heterozygous diploids as previously reported (*Nuckolls et al., 2017*). We grew these diploids to saturation in 5 mLs of rich YEL broth (0.5% yeast extract, 3% glucose, 250 mg/L of adenine, lysine, histidine, leucine, and uracil) overnight at 32°C. We then diluted 100 µL of these diploid cultures into 5 mLs of PM media (20 mLs of 50x EMM salts, 20 ml 0.4 M Na_2_HPO_4_, 25 mL 20% NH_4_Cl, 1 mL 1000x Vitamins, 100 µL 10,000x mineral stock solution, 3 g potassium hydrogen phthalate, 950 mL ddH_2_O, 25 mL of sterile 40% glucose after autoclaving, supplemented with 250 mg/L uracil) and again grew overnight at 32°C. The next day, we spun to pellet and resuspended the pellet in PM-N media (PM without NH_4_Cl). We shook the PM-N cultures for 4 h at 28°C. Then, we took 100 µL of the PM-N culture and mixed it with 100 µL of lectin (Sigma). We took 150 µl of this mixture and added it to a 35 mm glass bottom poly-D-lysine coated dish (MatTek corporation).

We waited five minutes to allow the cells to adhere. We then added 3 mLs of fresh PM-N to the dish (protocol modified from *Klutstein et al., 2015*). We imaged using a Zeiss Observer.Z1 wide-field microscope with a 63x (1.2 NA) oil-immersion objective and collected the emission onto a Hamamatsu ORCA Flash 4.0 using μManager software. We used BP 440-470 nm to excite GFP and collected BP 525-550 emission using a FT 495 dichroic, and mCherry with BP 530–585 nm excitation and LP 615 emission, using an FT 600 dichroic filter, acquiring images every 10 minutes.

For the Fluorescence Recovery After Photobleaching experiments, we imaged the cells on a Ti2-E (Nikon) microscope coupled to a CSU-W1 Spinning Disc (Yokogawa). GFP and mCherry were laser excited at 488 nm and 561 nm, respectively, through a 100x Plan Apochromat (Nikon, 1.45 NA) objective. The fluorescence emission of GFP was collected through a 525/36 nm filter, while the emission of mCherry was collected through a 605/25 nm filter. Both signals were collected on a Prime 95B camera (Photometrics). After an initial frame was recorded, the fluorescence of both fluorophores was bleached to background in a large field of view containing many asci, using 100% power from the 450nm, 550nm and 640nm laser lines. Once bleaching was completed, the recovery was recorded every 10 minutes for ∼6 hours total time. To generate the recovery curves, using Fiji, the average fluorescence intensity was quantified inside all spores containing GFP signal and all spores containing mCherry signal per frame, separately. Once the curves were obtained, all curves of the same fluorophore were normalized to the min and max intensity values and averaged to yield the final curves.

### Stress Experiments

For the stress experiments (S4 Fig), we generated saturated overnight cultures of haploid *Sp* strains [SZY4446 (Empty Vector), SZY4534 (*p^antidote^_mCh*)] in 5 mLs of rich YEL broth + the appropriate drug for selection of the construct. The next day, we took 1 mL of the saturated cultures and diluted into 4 mLs of fresh YEL+Drug. We did this three times to generate three separate diluted cultures. To one of the dilutions of each sample, we added tunicamycin to a final concentration of 0.5 μg/ml and shook at 32°C for one hour to induce ER stress. We placed the second dilutions at 40°C for one hour to induce heat stress. We placed the third dilution of each strain at 32°C as control samples. We imaged the samples using a Zeiss Observer.Z1 wide-field microscope with a 40x (1.4 NA) water objective and collected the emission onto a Hamamatsu ORCA Flash 4.0 using μManager software. We acquired the mCherry with BP 530–585 nm excitation and LP 615 emission, using an FT 600 dichroic filter. We quantified the raw intensity density of at least 50 cells per condition, after background subtraction. We repeated this experiment again on a different day to have an additional technical and biological replicate of each condition. The images for this experiment are shown at the same brightness and contrast for accurate comparison.

### Analysis of published Mei4 Chip-Seq data

FASTQ data (from *Alves-Rodrigues et al. 2016*) was retrieved from the European Nucleotide Archive (ENA), with accession number: ERP001894. Data was trimmed for quality using Trimmomatic (*Bolger et al., 2014*) and was aligned to *S. pombe* genome version ASM294v2 with bowtie2 (*Langmead and Salzberg, 2012*) using default parameters. Peaks were called using MACS2 with callpeak -gsize 1.38^e7^ and -q 0.05. Metagene plots were constructed by averaging the RPM (Reads Per Million) signal for individual genes [either the predicted *wtf* meiotic drive genes (*wtf4, wtf13,* top) or the *wtf* genes that are predicted to only encode antidote proteins (*wtf5, wtf9, wtf10, wtf16, wtf18, wtf20, wtf21, wtf25,* bottom)] across 400 bins and then averaging multiple genes to create a single profile followed by loess smoothing with a span of 0.5. For any reads that mapped to more than one location, only a single location, chosen at random, is reported.

## Acknowledgments

We would like to thank the members of the Zanders lab for their helpful comments on the paper. Original data underlying this manuscript can be accessed from the Stowers Original Data Repository at http://www.stowers.org/research/publications/libpb1562. This work was performed to fulfill, in part, requirements for NLN’s thesis research in the Graduate School of the Stowers Institute for Medical Research. This work was supported by The Stowers Institute for Medical Research (SEZ); the Searle Award (SEZ); National Institutes of Health (NIH) R00GM114436 and DP2GM132936 (SEZ); National Cancer Institute of the NIH under award number F99CA234523 (MABN); Eunice Kennedy Shriver National Institute of Child Health & Human Development of the NIH under Award Number F31HD097974 (NLN); The funders had no role in study design, data collection and analysis, or manuscript preparation. The content is solely the responsibility of the authors and does not necessarily represent the official views of the funders.

---

**S1 Figure.**
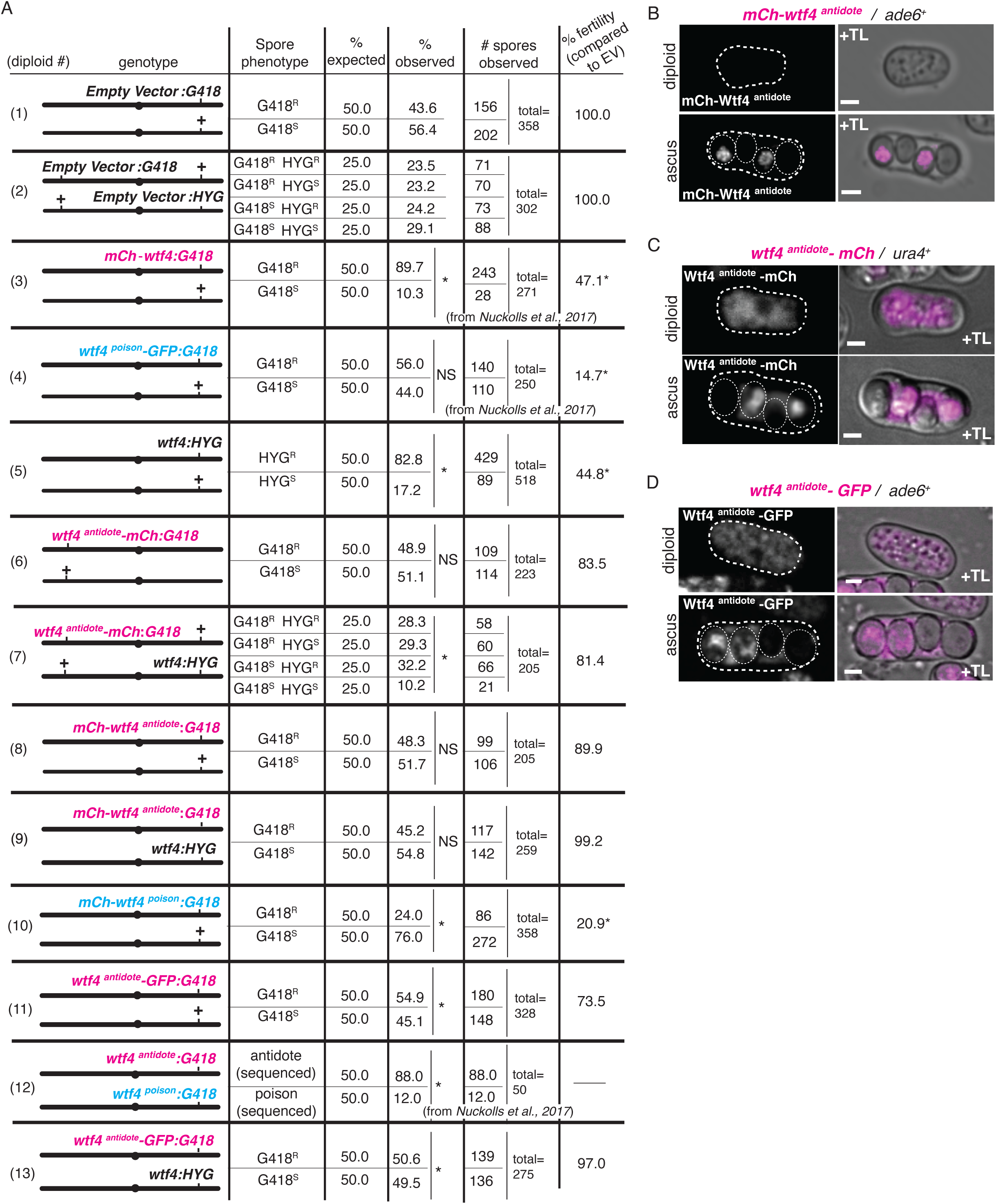
C-terminal tags reveal Wtf4^antidote^ expression prior to spore formation. (**A**) Allele transmission and fertility (assayed via viable spore yield) of 13 diploids with the depicted genotypes. The genotype column shows a cartoon depiction of the relevant genotype. The progeny phenotypes are then shown on the right. For diploids heterozygous at one locus (e.g. Diploid 1), two values are shown (top and bottom) that represent the two possible haploid genotypes. Spores exhibiting both parental phenotypes were considered diploid or aneuploid and were excluded from this table but can be found in S4 Data. The expected values assume Mendelian allele transmission. We used the viable spore yield assay (VSY) to quantify fertility with values normalized to the relevant empty vector control (*= p < 0.05, NS= not significant; G-test for allele transmission, Wilcoxon test for VSY, in comparison to the empty vector control). We compared diploids 3, 4, 5, 6, 8-13 to control diploid 1 and diploid 7 to control diploid 2. The data for diploids 1, 2, and 10-13 are also depicted in S4A Fig and the data for diploids 1 and 5 are also depicted in Fig 5C. The data from diploids 3, 4, and 12 were previously published in (*Nuckolls et al., 2017*). (**B**) Images of a heterozygous *mCherry-wtf4^antidote^/ade6+* diploid cell and mature ascus. mCherry-Wtf4^antidote^ is shown in magenta in merged images. (**C**) Images of a heterozygous *wtf4^antidote^-mCherry/ura4+* diploid cell and mature ascus. Wtf4^antidote^-mCherry is shown in magenta. Images of this cross are also presented in Fig 4C. (**D**) Images of a heterozygous *wtf4^antidote^-GFP/ade6+* diploid cell and mature ascus. Wtf4^antidote^-GFP is shown in magenta. Images of this cross are also presented in Fig 3A. All images were taken after 3 days on sporulation media. TL= transmitted light. All scale bars represent 2 μm. Not all images are shown at the same brightness and contrast to avoid over saturation of pixels in the brighter images.

**S2 Figure.**
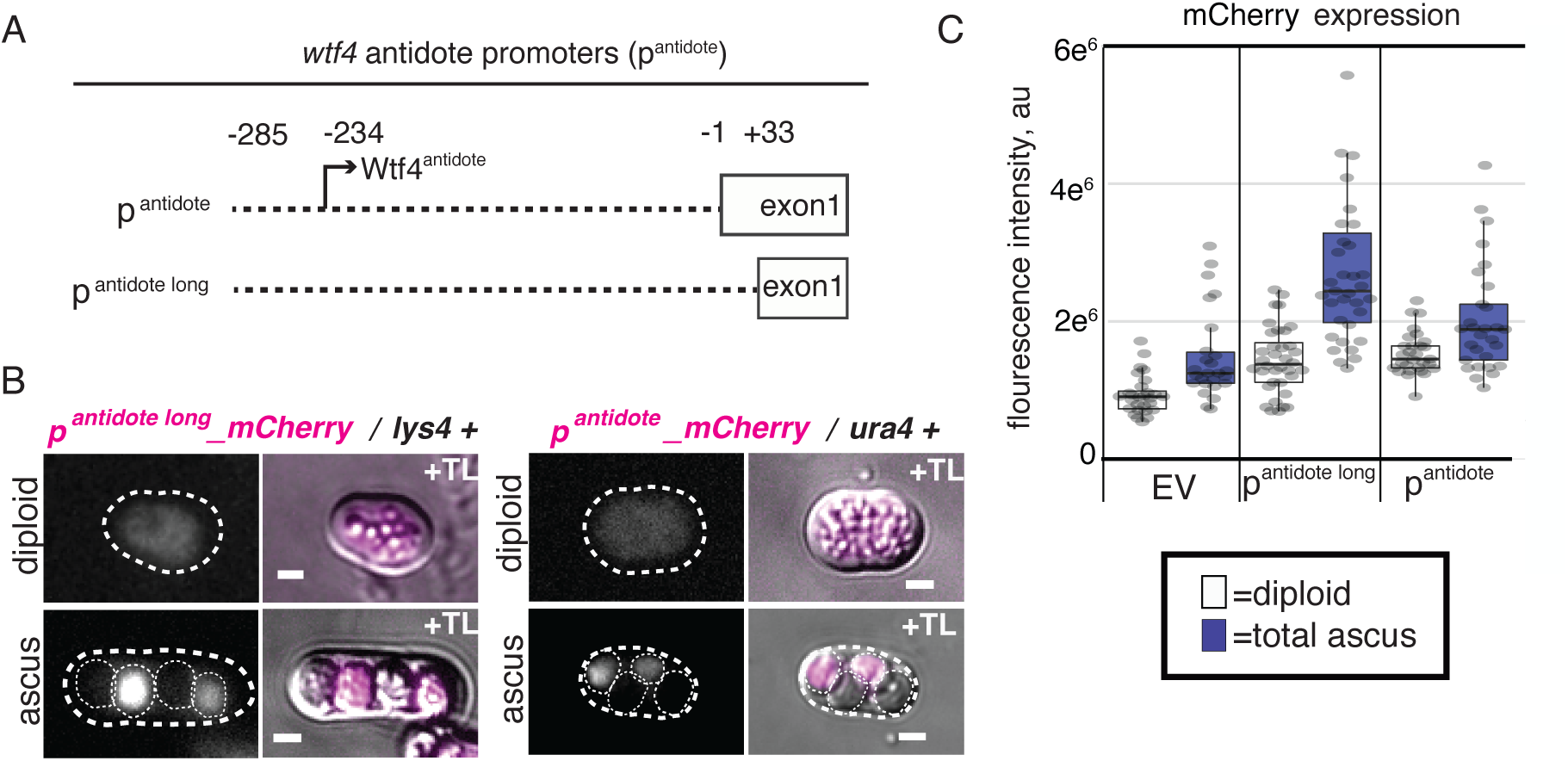
Expression of Wtf4^antidote^ promoters in diploids and asci. Depictions of (A) the two lengths of *wtf4^antidote^* promoters used in this study. (**B**) Images of the two different *p^antidote^*_*mCherry* reporter */ +* heterozygous diploids and asci. (**C**) Quantification of mCherry fluorescence within heterozygous (*p^antidote^*_*mCherry* reporter*/ +*) diploids and asci. At least 25 diploids and 25 asci were quantified per reporter. All images were acquired after 3 days on sporulation media. TL= transmitted light. All scale bars represent 2 μm. Images were taken at the same settings and are shown at the same brightness and contrast for accurate comparison.

**S3 Figure.**
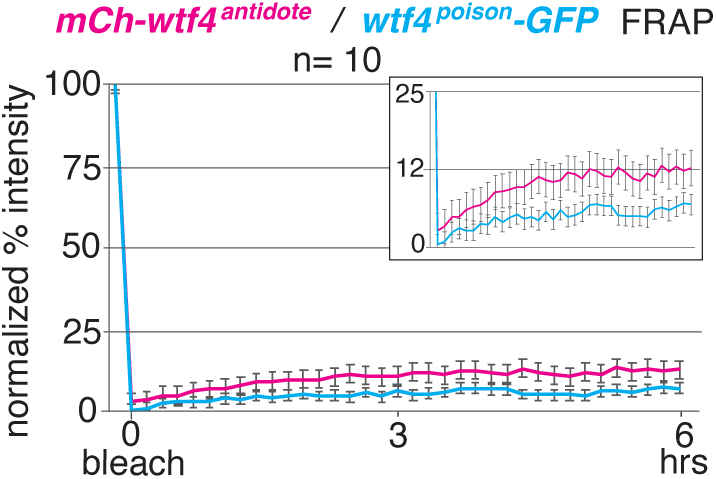
Both Wtf4 poison and antidote are produced at low level after spore individualization. <u>F</u>luorescence <u>R</u>ecovery <u>A</u>fter <u>P</u>hotobleaching (FRAP) of mature asci (n=10) generated from *mCherry-wtf4/ wtf4^poison^*-GFP diploids. Both mCherry (magenta line) and GFP (cyan line) were bleached to 0% intensity and recovery was quantified over 6 hours.

**S4 Figure.**
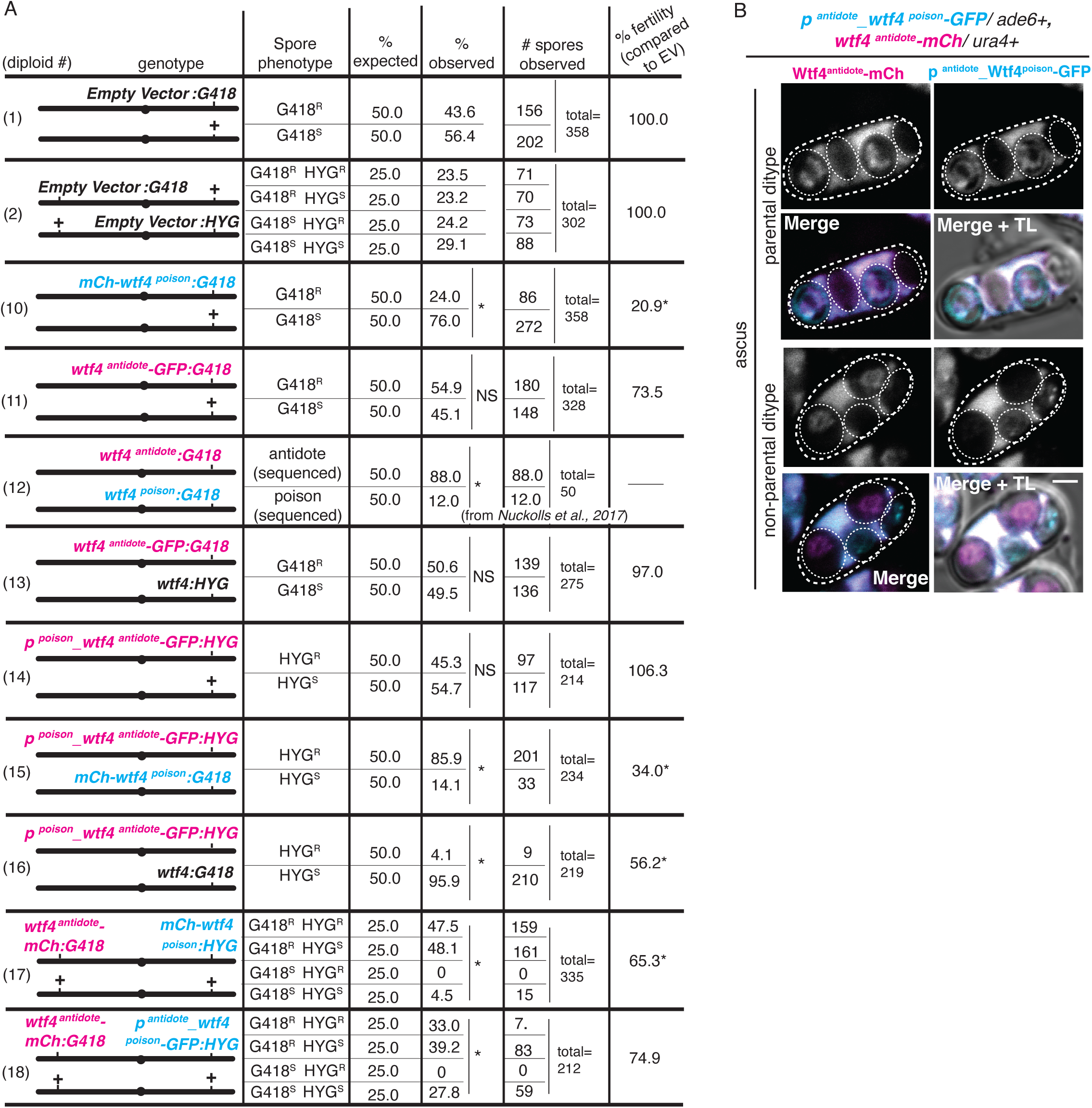
Wtf4^poison^ localization is altered when expressed from the p^antidote^ promoter. (**A**) Allele transmission and fertility (assayed via viable spore yield) of 11 diploids of the depicted genotypes. The genotype column shows a cartoon depiction of the relevant genotype. The progeny phenotypes are then shown on the right. For diploids heterozygous at one locus (e.g. diploid 1), two values are shown (top and bottom) that represent the two possible haploid genotypes. For diploids heterozygous at two loci, the loci used are unlinked and should segregate randomly. Spores exhibiting both parental phenotypes were considered diploid or aneuploid and were excluded from this table but can be found in S4 Data. The expected values assume Mendelian allele transmission. (*= p < 0.05, NS= not significant; G-test for allele transmission, Wilcoxon test for VSY, in comparison to the empty vector control). We compared diploids 10-16 to diploid 1 as the control and diploids 17 and 18 to diploid 2 as the control. The data for the control diploids 1 and 2 are also depicted in S1A Fig and Fig 5C, while diploids 10-13 are also depicted in S1A Fig. (**B**) Images of a *p^antidote^*_*wtf4^poison^-GFP/ade6+, wtf4^antidote^- mCherry/ura4+* parental and non-parental ditype asci. p^antidote^_Wtf4^poison^-GFP is shown in cyan and Wtf4^antidote^-mCherry is shown in magenta in merged images. TL= transmitted light. All scale bars represent 2 μm. All images acquired after 3 days on sporulation media. Not all images are shown at the same brightness and contrast to avoid over saturation of pixels in the brighter images.

**S5 Figure.**
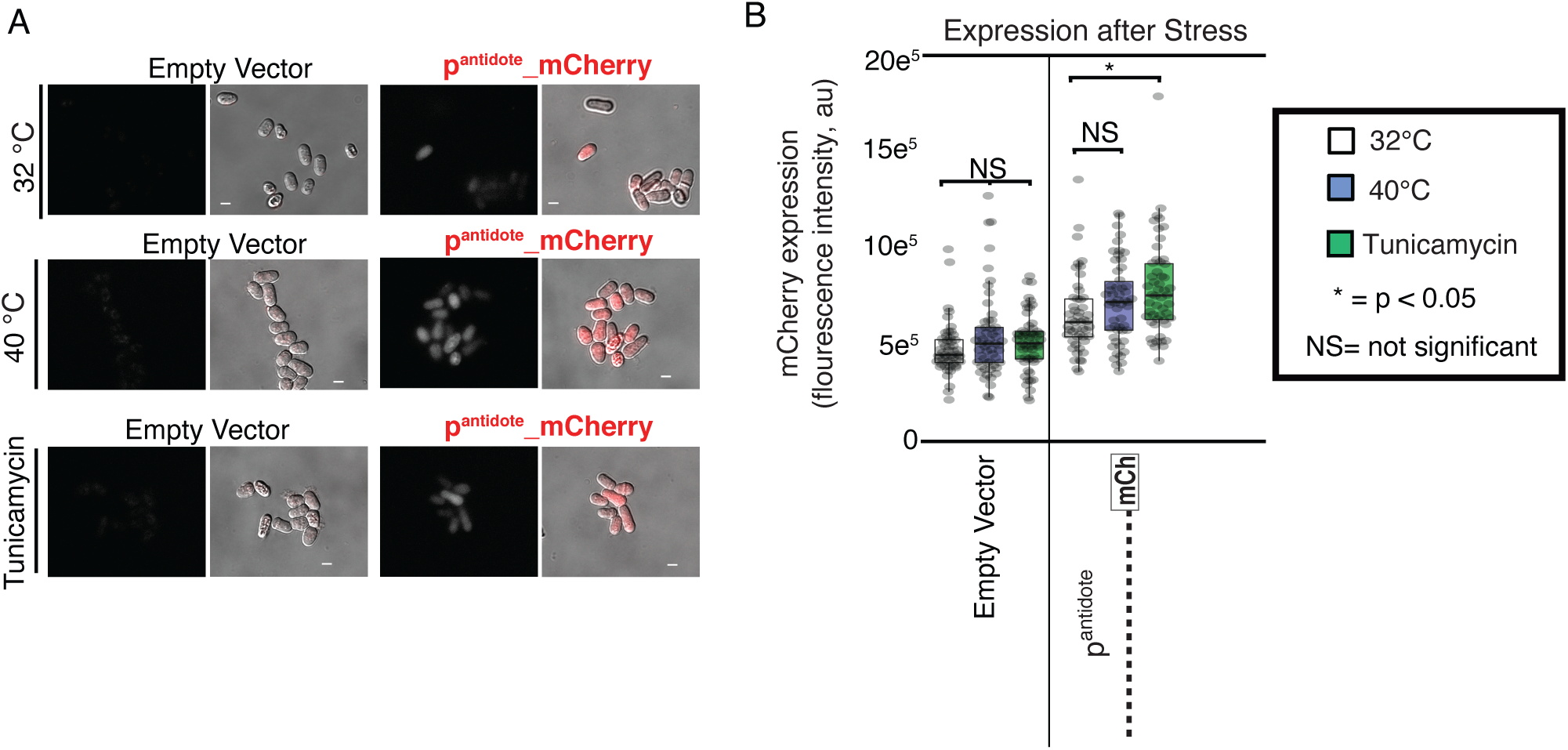
p^antidote^ promoter promotes expression after ER stress. (**A**) Images of haploid *S. pombe* cells carrying either an empty vector or the *p^antidote^-mCherry* reporter construct that were grown for an hour at 32°C (control), 40°C (heat-shock), or at 32°C with 0.5 µL/mL Tunicamycin (to induce endoplasmic reticulum stress). Scale bars represent 4 µM. These experiments were all completed at the same time to ensure accurate comparison and are shown at normalized brightness and contrast. We also repeated the analyses a second time to ensure adequate replicates were done (see methods). (**B**) The quantification of the samples shown in part A. (*= p < 0.05, NS= not significant, t-test). n>50 cells for all conditions.

**S1 Data.**
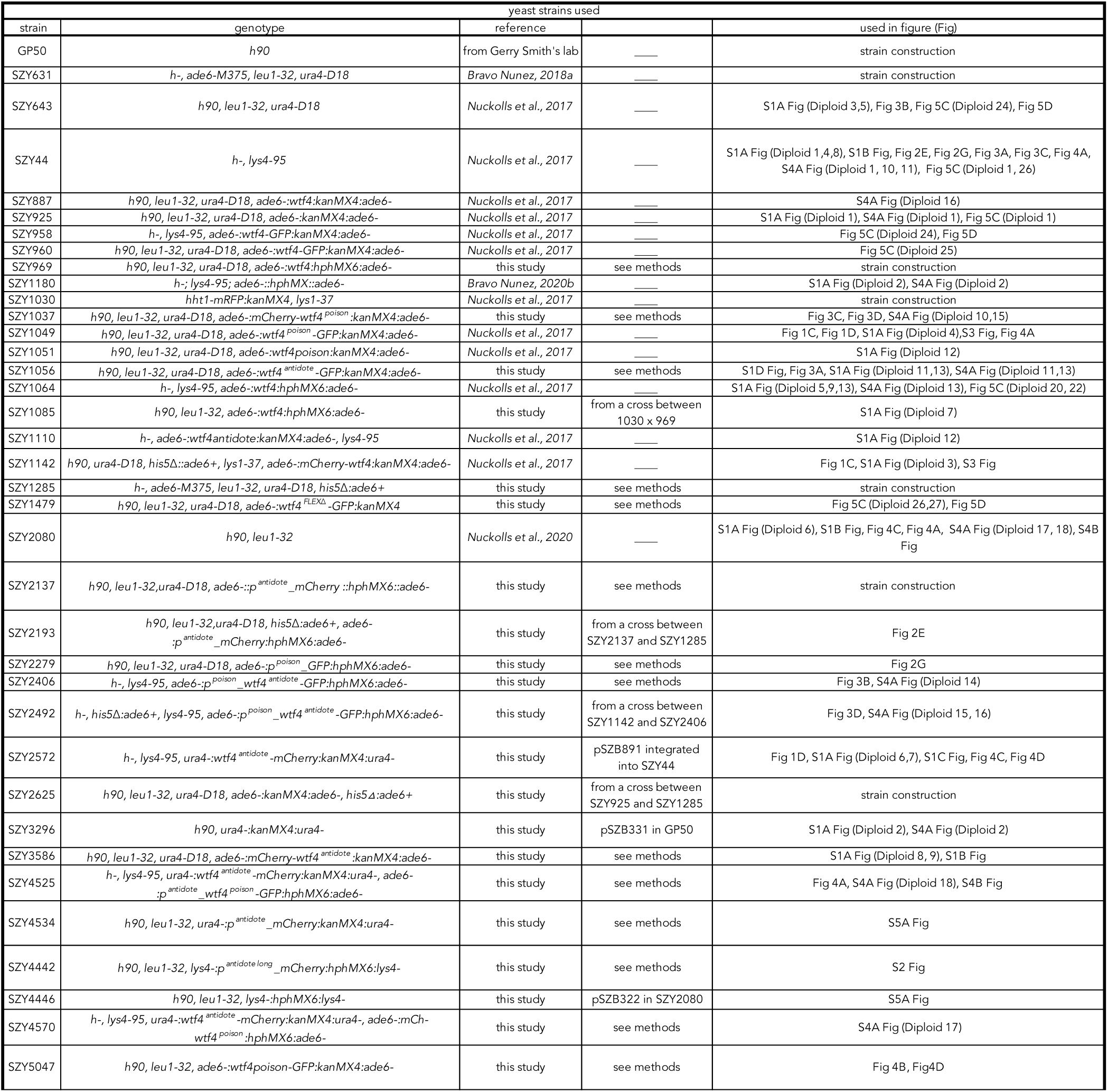
Yeast strains used. Column 1 is the name of strain used, while column 2 refers to the genotype. Columns 3 lists the reference for the yeast strain. If it was made in this study, we also detail how the strain was made in column 4. Column 5 lists the figure(s) in which the strain was used.

**S2 Data.**
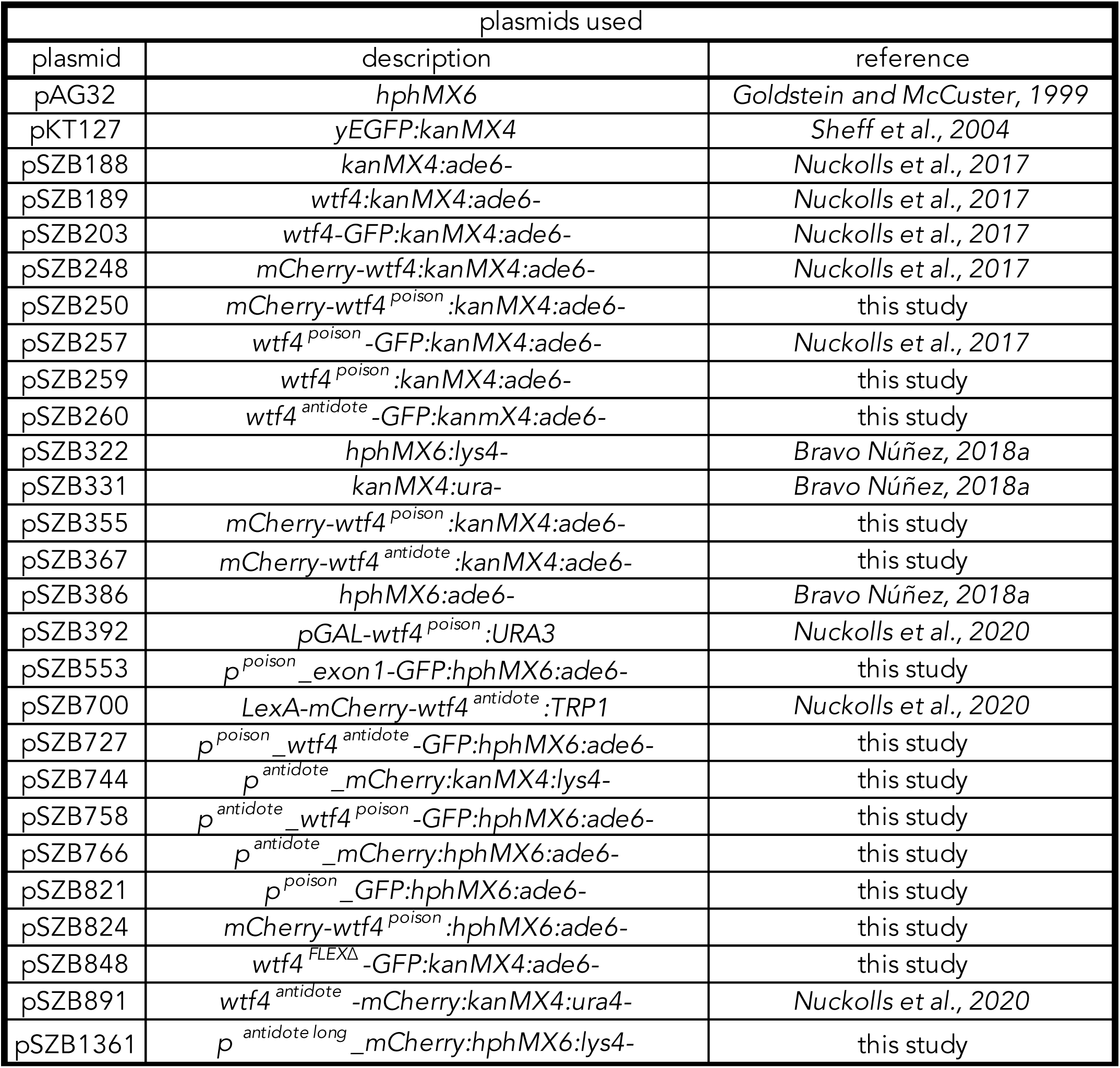
Plasmids used. Column 1 is the name of plasmid used. Column 2 gives a short description of the plasmid. Columns 3 lists the reference for the plasmid.

**S3 Data.**
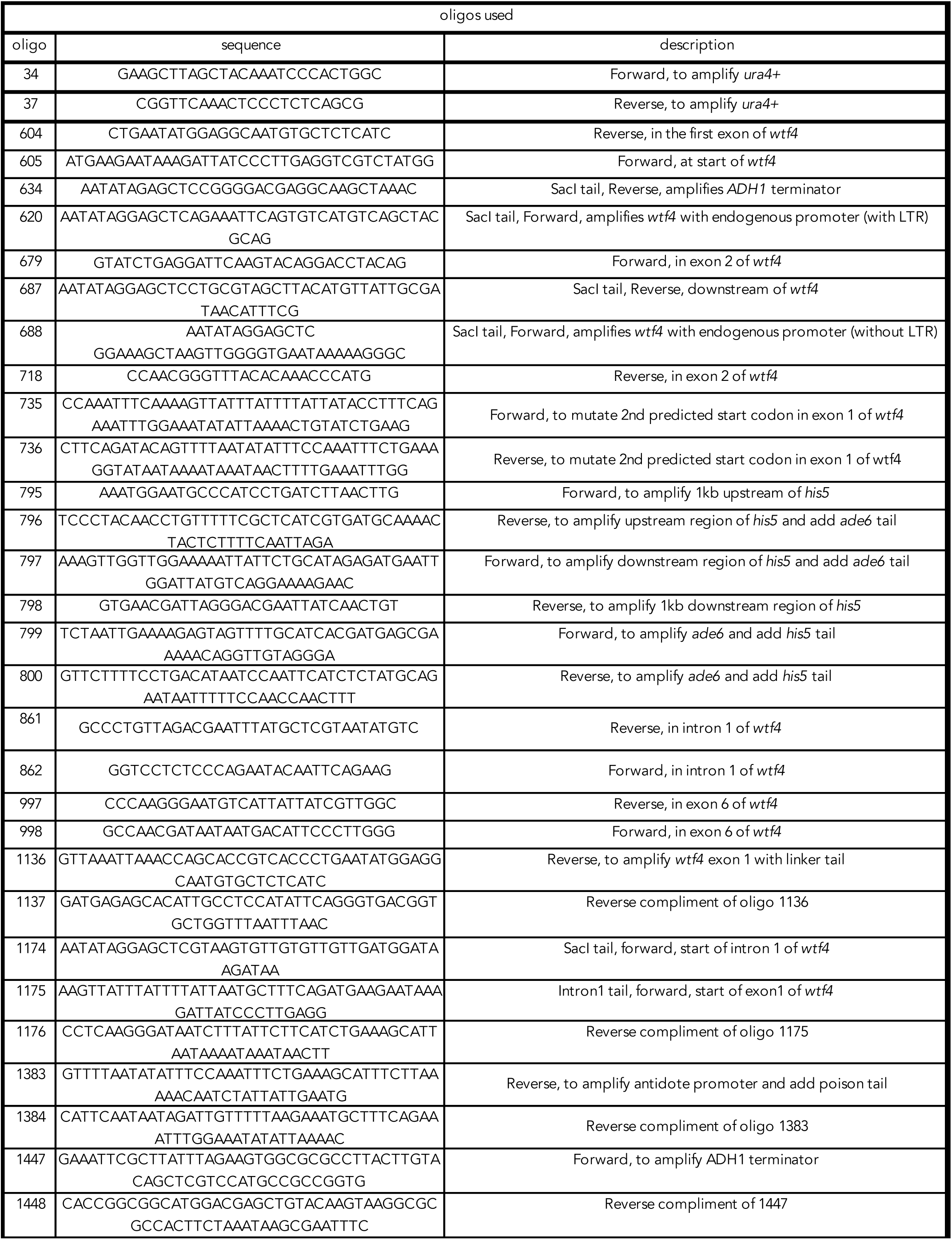
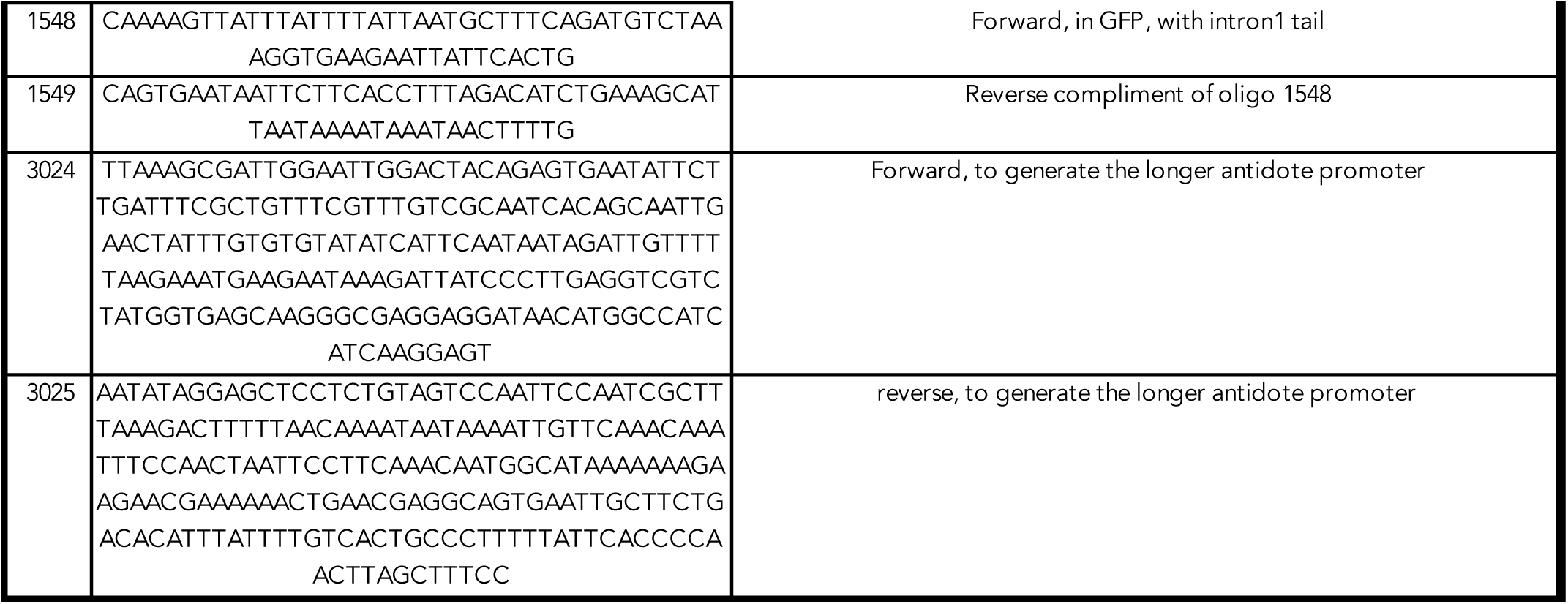
Oligos Used. Column 1 is the name of oligo used. Column 2 details the sequence of the oligo, while column 3 gives a short description.

**<u>S4 Data.</u> Genetics and Viable Spore Yield (VSY) data for all diploids presented.** Tab 1 is a summary of all the allele transmission and VSY data for all 22 diploids. Each subsequent tab contains the raw data for individual crosses. The tab name contains the diploid number, corresponding to the diploid number in the paper, and the two strains crossed to generate the diploid. The top table in each tab lists the VSY data, including the data for at least 3 independent diploids (A-X) per cross. The number of colonies for the diploid dilutions and the corresponding spore dilutions are listed, as well as the average viable spore yield, standard deviation, and the p-value when compared to the appropriate control (Wilcoxon test, *= p < 0.05, NS= not significant). The bottom table contains the allele transmission data. We detail the genotypes of the two strains crossed to generate the diploid (allele 1 and allele 2) and the transmission frequencies of the two alleles, as well as a control locus. We show the data with and without disomes in the table, but used the data excluding disomes in the figures. We also list the p-values compared to appropriate control (G-test, *= p < 0.05, NS= not significant).

Provided as an excel sheet.

**<u>S5 Data.</u> Raw data of image quantification.** Each tab contains the raw data of quantifications presented in the manuscript. The first tab contains the quantifications depicted in Figure 2F and 2H of reporter (promoter-FP) expression in spores. For each ascus, we detail the mean fluorescence intensity of the 4 individual spores and the total of the four spores. We then divided each individual spore by the total to get the % of total spore fluorescence inside a given spore. The second tab details the quantification depicted in Figure 3D of % intensity inside spores of given fluorescent proteins. We show the % of fluorescent protein inside spores in three different genotypes (*p^poison^_wtf4^antidote^-GFP/mCh-wtf4^poison^, p^poison^_wtf4^antidote^-GFP/ade6+,mCh-wtf4^poison^/ade6+).* The third tab details the quantifications shown in S2C Figure of fluorescence intensity inside asci and diploids for the following constructs: p^antidote^_mCherry, p^antidote long^_mCherry, and the empty vector control. The fourth tab details the Fluorescence Recovery After Photobleaching (FRAP) quantifications depicted in S3 figure. We normalized the fluorescence intensity before bleaching to 1. Once bleaching was completed, the recovery was recorded every 10 minutes for approximately 6 hours. The fluorescence levels of both mCherry-Wtf4 and Wtf4^poison^-GFP are recorded for each time point. The fifth tab details Figure 4E, the % intensity inside spores of given fluorescent proteins. We show the % of Wtf4^poison^-GFP inside spores in three different genotypes: *p^antidote^-wtf4^poison^-GFP/ade6+, wtf4^antidote^-mCh/ura4+; wtf4^poison^-GFP/ade6+*; and *wtf4^poison^-GFP/ade6+, wtf4^antidote-^mCh/ura4+.* The final tab details the quantifications shown in S5B Figure of fluorescence intensity of p^antidote^_mCherry in cells with and without stress.

Provided as an excel sheet

